# Optogenetic control of small GTPases reveals RhoA-mediated intracellular calcium signaling

**DOI:** 10.1101/2020.07.13.200089

**Authors:** Hironori Inaba, Qianqian Miao, Takao Nakata

**Author notes:** Corresponding author: Takao Nakata.

## Abstract

Rho/Ras family small GTPases are known to regulate numerous cellular processes, including cytoskeletal reorganization, cell proliferation, and cell differentiation. These processes are also controlled by Ca^2+^, and consequently, crosstalk between these signals is considered likely. However, systematic quantitative evaluation is not yet reported. Thus, we constructed optogenetic tools to control the activity of small GTPases (RhoA, Rac1, Cdc42, Ras, Rap, and Ral) using an improved light-inducible dimer system (iLID). Using these optogenetic tools, we investigated calcium mobilization immediately after small GTPase activation. Unexpectedly, we found that a transient intracellular calcium elevation was specifically induced by RhoA activation in RPE1 and HeLa cells. These findings were confirmed in all the cell lines tested in this study. Of note, molecular mechanisms were identified to be different among cell types. In RPE1 and HeLa cells, RhoA activated phospholipase C epsilon (PLCε) at the plasma membrane, which in turn induced Ca^2+^ release from the endoplasmic reticulum (ER). The RhoA-PLCε axis induced calcium-dependent NFAT nuclear translocation, suggesting it does activate intracellular calcium signaling.

## Introduction

Small GTPases of the Ras superfamily have been identified as molecular switches because they exist in two states, namely a GTP-bound state (“ON”) and a GDP-bound state (“OFF”) (1, 2). These states are known to be regulated by guanine nucleotide exchange factors (GEFs) as activators and GTPase-activating proteins (GAPs) as inactivators. Rho and Ras subfamily small GTPases localize to the plasma membrane, respond to extracellular stimuli, and regulate a variety of biological processes, including cytoskeletal reorganization, cell proliferation, and cell differentiation (1, 2). Several of these biological processes are also regulated by the universal second messenger ionic calcium (Ca^2+^) (3–5). Thus, functional links must exist between the signaling pathways regulated by Rho/Ras family small GTPases and Ca^2+^. Small GTPases and Ca^2+^ share some downstream factors that are coordinately regulated. For example, RhoA and Ca^2+^ regulate myosin II activity via myosin light chain (MLC) phosphorylation (6, 7), and Ras and Ca^2+^ coordinate the extracellular signal-regulated kinase (ERK)/mitogen-activated kinase (MAPK) signaling pathway (8, 9). In addition, small GTPases and Ca^2+^ are known to regulate each other’s functions. Specifically, many GEFs and GAPs are regulated both positively and negatively by Ca^2+^ (4, 10), and some small GTPases regulate intracellular calcium signaling by activating phospholipase C (PLC) (11, 12). PLC converts phosphatidylinositol 4,5-bisphosphate [PI(4,5)P_2_] to two second messengers: diacylglycerol (DAG) and inositol trisphosphate (IP_3_). IP_3_ reportedly binds to the IP_3_ receptor (IP3R) to release Ca^2+^ from the endoplasmic reticulum (ER). This PLC-mediated calcium influx is the major calcium signaling pathway in non-excitable cells.

Despite the importance of crosstalk between small GTPases and intracellular calcium, details of these processes remain poorly understood. In particular, assessment of the influence of small GTPases on intracellular calcium concentrations immediately after activation has been difficult because this activity cannot be directly controlled in cells. However, optogenetics has changed this situation over the last decade.

Optogenetics is a pivotal tool for advancing cell biology because it enables the control of specific signaling molecules at high spatiotemporal resolution both *in vitro* and *in vivo* (13–15). The optogenetic control of small GTPases was first reported by Hahn’s group (16). In their study, constitutively active mutants of Rac1 and Cdc42 were fused to the blue light-excited light-oxygen-voltage-sensing domain 2 (LOV2) of phototropin from *Avena sativa* (17). Photoactivatable (PA)–Rac1 and PA–Cdc42 were inactive in the dark because of steric hindrance of effector binding sites by the LOV2 domain. Blue light irradiation induces conformation changes in the alpha helix (Jα) that connects LOV2 domains to small GTPases, allowing them to bind effectors. However, this approach was difficult to optimize between “ON” and “OFF” states for other small GTPases. Therefore, the plasma membrane translocation of their specific GEFs with light-induced heterodimeric systems, such as CRY2-CIBN (18), iLID (19), TULIP (20) and PhyB-PIF (21) systems, has been broadly used to regulate the activity of small GTPases including Rac1 (19, 21, 22), Cdc42 (19, 21, 22), RhoA (23–25), Ras (26), and Ral (27).

We have constructed optogenetic tools to control the activity of six members of the Rho and Ras subfamily GTPases (RhoA, Rac1, Cdc42, Ras, Rap, and Ral) by light-inducing GEF translocation to the plasma membrane using the iLID system. Using these optogenetic tools, we examined small GTPase-mediated intracellular calcium mobilization for the first time. Unexpectedly, transient elevation of intracellular calcium concentrations was only induced by optogenetic RhoA activation. These RhoA-mediated calcium transients were observed in all cell types examined, but the molecular mechanisms were different among the cell types. Furthermore, we found that RhoA activated PLCε in RPE1 and HeLa cells, which induced intracellular calcium signaling.

## Results

### Construction of optogenetic tools for controlling small GTPase activity

Specific control of Rho/Ras family small GTPase activity at high spatiotemporal resolution was achieved using optogenetic tools. Among the several light-inducible heterodimerization systems, we chose the iLID system because of the reasons that follow: (1) it is based on the *As*LOV2 domain that can work without exogenously adding a chromophore to mammalian cells; (2) iLID-SspB heterodimerization can be controlled by blue-light with rapid on/off kinetics (seconds), which is suitable in controlling small GTPase activity at high spatiotemporal resolution; and (3) the molecular weight of proteins is small, allowing high-level expression in cells and relative ease of establishing a lentivirus vector for stable cell lines.

To evaluate the system, we first constructed an optogenetic tool to control RhoA activity (opto-RhoA; **Figure 1A**). Similar to previous studies (23–25), the DH domain of the RhoA-specific GEF LARG (LARG-DH) was fused to SspB and the fluorescent protein mVenus. The CAAX motif of K-Ras (CAAX^Kras^) was fused to the C-terminus of iLID to induce its localization to the plasma membrane. Further, a nuclear export signal (NES) was inserted between mVenus and SspB to avoid nuclear co-localization, as originally reported (19). Cells were transfected with the single plasmid mVenus-SspB-LARG-DH, and iLID-CAAX^KRas^ was linked with the cDNA of the self-cleaved P2A peptide. Therefore, the observation of mVenus fluorescence confirmed that both fragments were expressed. In the dark, mVenus-SspB-LARG-DH was diffusely localized in the cytoplasm (**Figure 1B, 458-nm light OFF**). Blue light irradiation using a 458-nm laser induced iLID–SspB heterodimerization, which in turn induced the localization of LARG-DH to the plasma membrane, at which it could activate RhoA (**Figure 1B, 458-nm light ON**).

**Figure 1.**
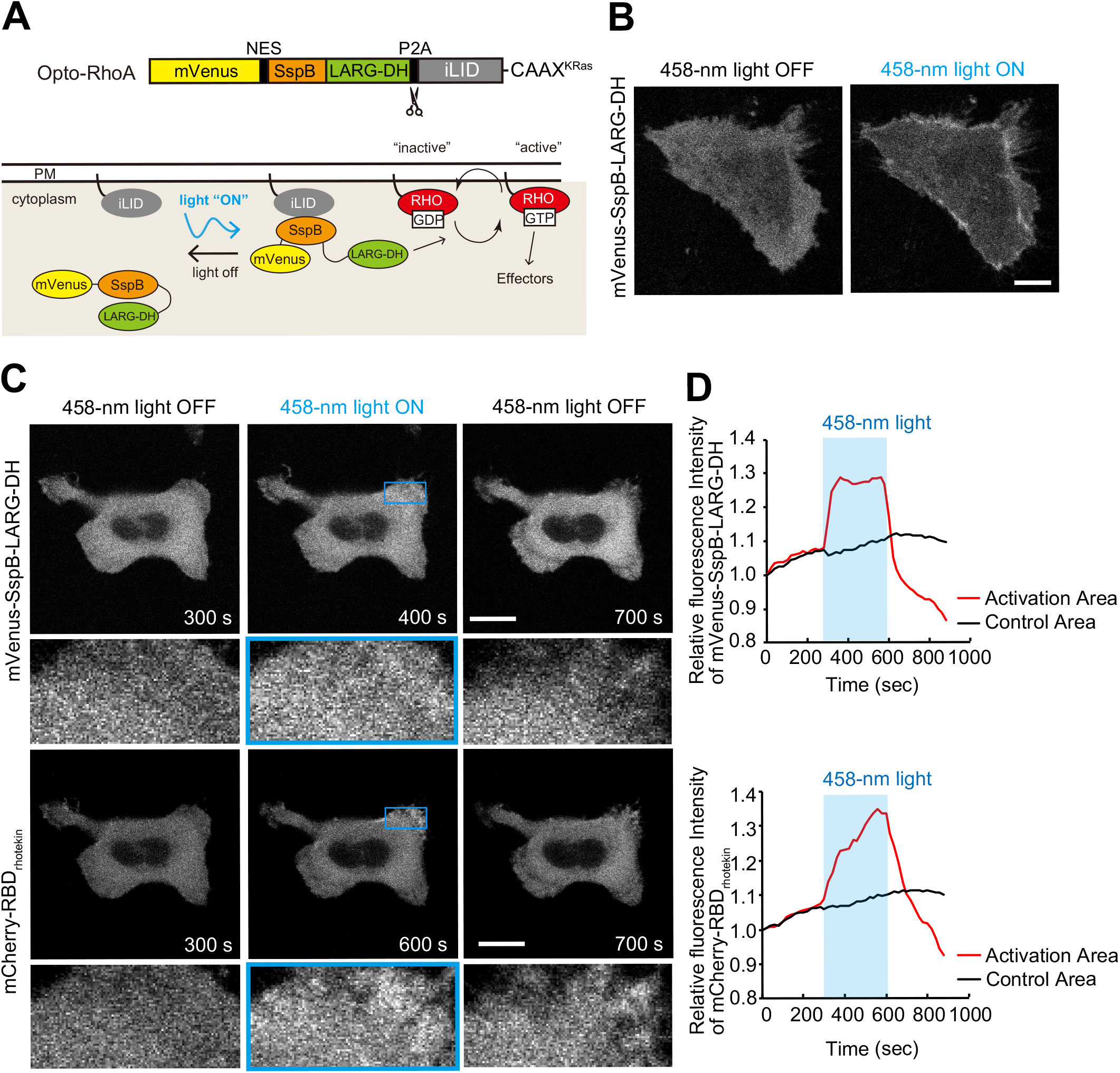
Construction of an optogenetic tool to control RhoA activity. **(A)** Schematic of opto-RhoA. Spatiotemporal control of RhoA activity is achieved using an optogenetic probe to recruit RhoA-specific GEF LARG to the plasma membrane. An iLID molecule is anchored to the plasma membrane via the CAAX motif, while a protein consisting of SspB fused to the DH domain of LARG is distributed throughout the cytosol. When irradiated with blue-light, iLID undergoes a conformational change exposing a binding site for SspB, and driving LARG-DH to the plasma membrane, where it activates RhoA. **(B)** Representative images of HeLa cells expressing opto-RhoA before (458-nm light OFF) and after (458-nm light ON) light irradiation with a 458-nm laser. Scale bar = 20 μm. **(C)** Representative images of HeLa cells expressing opto-RhoA and mCherry-RBD_rhotekin_. Opto-RhoA was locally activated by a 458-nm laser within 300–600 s every 20 s. Blue boxes in images indicate the activated area, and lower panels showed high magnification images for this area. (**D**) Quantification of local intensity increases of mVenus-SspB-LARG-DH (**top**) and mCherry-RBD_rhotekin_ (**bottom**) from the images shown in **(C)**. The entire area of the cell excluding the region of activation is indicated as the control area. Activation period is indicated by a blue background. Experiments were repeated seven times.

Then, we characterized this light-induced translocation using the mCherry version of opto-RhoA, in which mVenus was replaced with mCherry (**Supplementary Figure S1**). Blue light was irradiated using a laser line of confocal microscope, while imaging CFP (458-nm), GFP (488-nm) and, YFP (515-nm) channels. Translocation occurred in a laser power-dependent manner following irradiation with both 458- and 488-nm-light, and a laser power of 2% was sufficient for maximum translocation (**Supplementary Figure S1A–B**). Upon light irradiation, mCherry-SspB-LARG-DH rapidly translocated to the plasma membrane. After irradiation was terminated, the localization reverted within 1 min similarly as observed in the original iLID system (**Supplementary Figure S1D**) (19). Conversely, 515-nm irradiation at 1–2% laser power did not affect the localization of mCherry-SspB-LARG-DH (**Supplementary Figure S1C**). The use of 515-nm light with higher laser power (5 and 10%) of 515-nm light affected the localization in a laser power-dependent manner.

Next, we examined RhoA activation by opto-RhoA using a RhoA activity reporter, namely mCherry fused with the Rho-binding domain derived from the RhoA-specific effector rhotekin (mCherry-RBD_rhotekin_) (28). Opto-RhoA and mCherry-RBD_rhotekin_ were transfected into HeLa cells, and cells were locally irradiated with a 458-nm laser (**Figure 1C and Movie 1**). During irradiation, mVenus-SspB-LARG-DH rapidly accumulated in the irradiated area, whereas mCherry-RBD_rhotekin_ accumulated gradually (**Figure 1D**). Their accumulation disappeared after irradiation was terminated. These data suggest that RhoA activity can be controlled spatiotemporally using opto-RhoA as previously reported (23–25).

We next constructed optogenetic tools for five other members of Rho and Ras family GTPases (Rac1, Cdc42, Ras, Rap, and Ral) implicated in regulating intracellular calcium signaling (11, 12), as well as empty control using the same system (**Figure 2A**). LARG-DH of opto-RhoA was replaced with the GEF domains of specific GEFs for each small GTPase (see details in the Experimental procedure section). Most of these constructs were previously applied in optogenetic analyses (19, 26, 27). We named these optogenetic tools opto-X (X: target GTPase).

**Figure 2.**
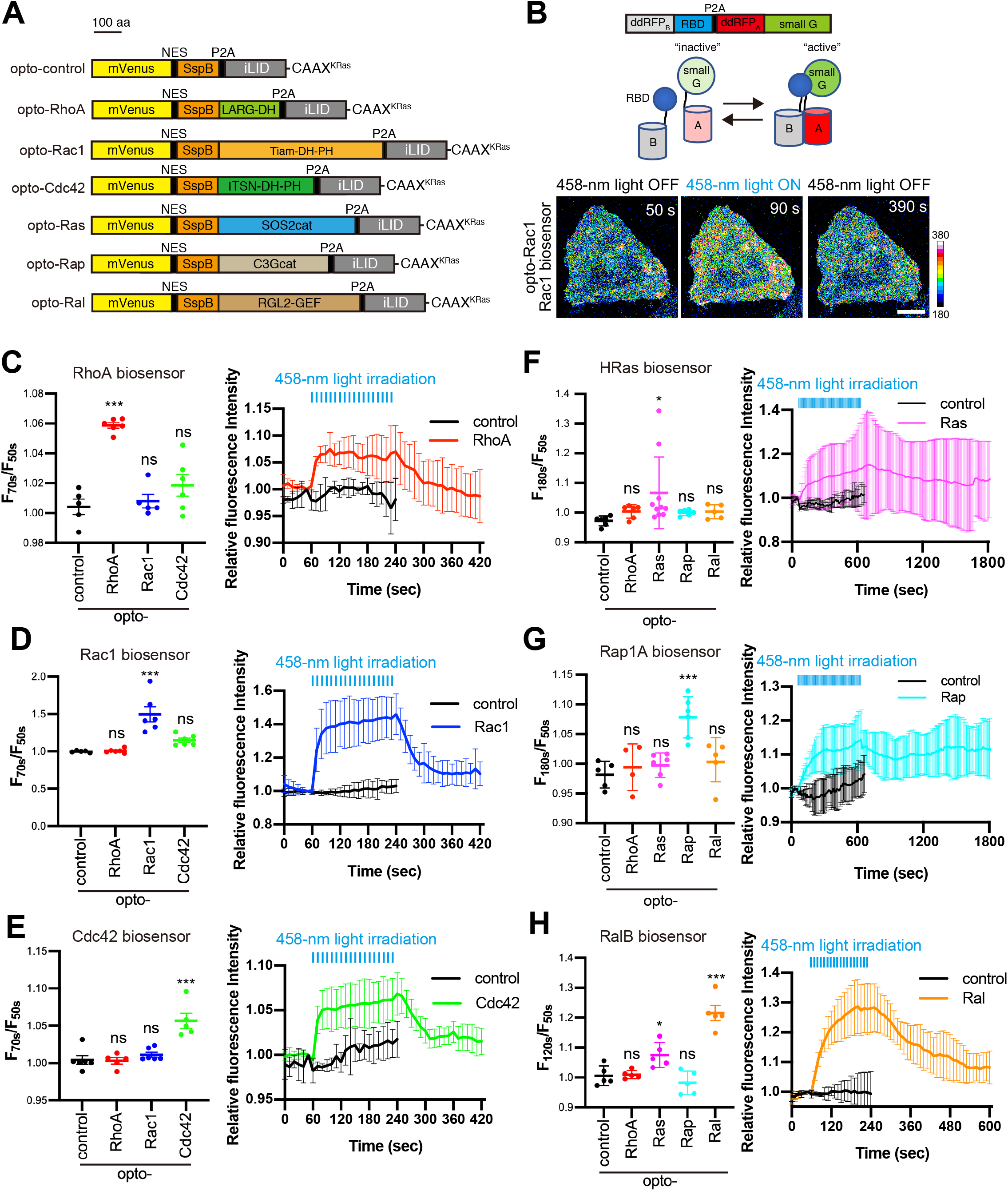
Construction and characterization of optogenetic tools to control small GTPase activity. (**A**) Schematic of opto-GTPases. (**B**) Schematic of ddRFP-based small GTPase sensor (top and middle). Example images for the experiments (bottom). HeLa cells transiently expressing opto-Rac1 and Rac1 biosensors were irradiated using a 458-nm laser every 10 s during a period of 60–230 s. During irradiation, an increase in the fluorescence intensity of the Rac1 biosensor indicated an increase of Rac1 activity. The color bar indicates the range of Rac1 biosensor intensity. Scale bar = 20 μm. (**C–H**) HeLa cells transiently expressing opto-GTPases and small GTPase biosensors were observed using confocal microscopy in Leibovitz’s L-15 medium. Opto-GTPases were activated by a multi-argon 458-nm laser every 10 s during a period of 60–230 s (**C–E, H**) or 60–350 s (**F–G**). Changes in the fluorescence intensity of biosensors before and after irradiation with the 458-nm light (**left**). Time course of small GTPase activity in cells expressing opto-control and opto-GTPase corresponding to each biosensor (**right**). Data are presented as the means ± SD. More than five cells were analyzed for each condition, and the actual number of cells is presented as the number of data points. Time course for all opto-GTPases is presented in Supplementary Figure S3. Data were analyzed by one-way ANOVA followed by Dunnett’s test (**C–E** and **G–H**) or Brown-Forsythe ANOVA followed by Dunnett’s T3 test (**F**) between cells expressing opto-control and other groups. ANOVA *F* = 24.5 and *p* < 0.0001 (**C**), *F* = 17.75 and *p* < 0.0001 (**D**), *F* = 15.58 and *p* < 0.0001 (**E**), *F* = 3.571 and *p* = 0.0417 (**F**), *F* = 8.750 and *p* = 0.0003 (**G**), *F* = 29.13 and *p* < 0.0001 (**H**). ***, *p* < 0.001; **, *p* < 0.01, *, *p* < 0.05; and ns, not significant; Dunnett’s test.

To evaluate these optogenetic tools, we first assessed their light-induced translocation using mCherry versions of opto-GTPases (**Supplementary Figure S2**). Excluding opto-Rap, the constructs exhibited similar translocation kinetics both during and after blue light irradiation even though they exhibited different levels of translocation. Opto-Rap translocation was slower than other opto-GTPases, and it took approximately 50 s to reach maximum translocation.

Next, to evaluate the functions of these tools, we employed genetically encoded red fluorescence intensity-based small GTPase biosensors (29) (**Figure 2B**). The biosensors consisted of a dimerization-dependent red fluorescent protein (ddRFP)_B_-fused Rho/Ras binding domain (RBD) and ddRFP_A_-fused small GTPases. When small GTPases were activated, their specific RBD interacted with small GTPases. Then, ddRFP_A_ and ddRFP_B_ formed a dimer that in turn increased their fluorescence intensity. Rac1, Cdc42, and HRas biosensors were already reported (29). Hence, we constructed RhoA, Rap1A, and RalB biosensors in this study. RBDs were selected to align with the FRET biosensors of these small GTPases, and/or baits for active GTPases GST-pulldown assay as follows: RBD derived from rhotekin for RhoA, RalGDS for Rap1A, and Sec5 for RalB. RhoA biosensor exhibited an increase in fluorescence intensity upon treatment of lysophosphatidic acid (LPA), and Rap1 and RalB biosensors exhibited that the same upon treatment of epidermal growth factor (EGF), thus confirming these biosensors can monitor GTPases activities (data not shown). As expected, fluorescence intensity was significantly increased after blue-light illumination in cells co-expressing opto-GTPases and the corresponding small GTPase biosensors (**Figure 2C–H and Supplementary Figure S3**). The changes in fluorescence intensity of H-Ras biosensor by opto-Ras were highly variable and the *p*-value was higher than others. Therefore, to evaluate the function of opto-Ras, we also performed ERK nuclear translocation assay, which is generally used to evaluate Ras activity (26, 30) (**Supplementary Figure S4**). Opto-Ras induced ERK nuclear translocation significantly, although the variability was also large. This experiment confirmed that opto-Ras activates Ras as intended. Interestingly, both the activation and inactivation of Rho family small GTPases (Rho, Rac1, and Cdc42) occurred with similar speed as opto-GTPase translocation (**Figure 2C–E**). By contrast, the activation and inactivation of Ras family small GTPases (HRas, Rap1A, and RalB) were more gradual than those of Rho family small GTPases (**Figure 2F–H**). The activities of both HRas and Rap1A persisted even 20 min after the irradiation was terminated. We also examined the specificity of these tools (**Figure 2C–H and Supplementary Figures S3 and S4**) regardless of whether they activate other family members of small GTPases, especially for the same family members. Among these optogenetic tools, opto-Cdc42 activated both RhoA and Rac1 (**Figures 2C–D and Supplementary Figure S3A–B**), although not in a statistically significant manner, and opto-Ras activated RalB (**Figure 2H and Supplementary Figure S3F**). The activation for Rac1 by opto-Cdc42 was weaker than that by opto-Rac1. In addition, the activation of RhoA by opto-Cdc42 was more gradual than by that by opto-RhoA. Meanwhile, the optogenetic activation of both Rac1 and RhoA by Cdc42 has been reported (22, 31). Therefore, these results likely reflect indirect effects via Cdc42 activation rather than a specific process (see also the Discussion section). Conversely, RalB activation by opto-Ras was not surprising because Ral is a well-known downstream factor of Ras (2). In summary, all opto-GTPases could specifically activate their corresponding GTPases as intended.

### Screening using opto-GTPases for induction of changes in intracellular calcium concentrations

We monitored changes of intracellular calcium concentrations with a genetically encoded red fluorescent calcium indicator R-GECO1 that is known to detect physiological calcium changes as found during neural activity and muscle contraction (32). Opto-GTPases and the calcium reporter R-GECO1 were transfected into human non-transformed (RPE1) and cancer (HeLa) cells. Changes of intracellular calcium concentrations induced by persistent activation of small GTPases were observed via confocal microscopy. Among the GTPases, only RhoA was identified to induce transient elevation of intracellular calcium concentrations both in RPE1 and HeLa cells **(Figure 3, Supplementary Figure S5, and Movie 2)**. We also tested a YA mutant of opto-RhoA (opto-RhoA^YA^) that possessed a LARG Y940A mutation that abolished GEF activity (33). Opto-RhoA^YA^ rarely induced calcium transients (1 out of 136 RPE1 cells and 0 out of 133 HeLa cells; **Figure 3C**), confirming that calcium transients were induced by RhoA activation and not by blue-light irradiation or by photoswitch translocation to the plasma membrane.

**Figure 3.**
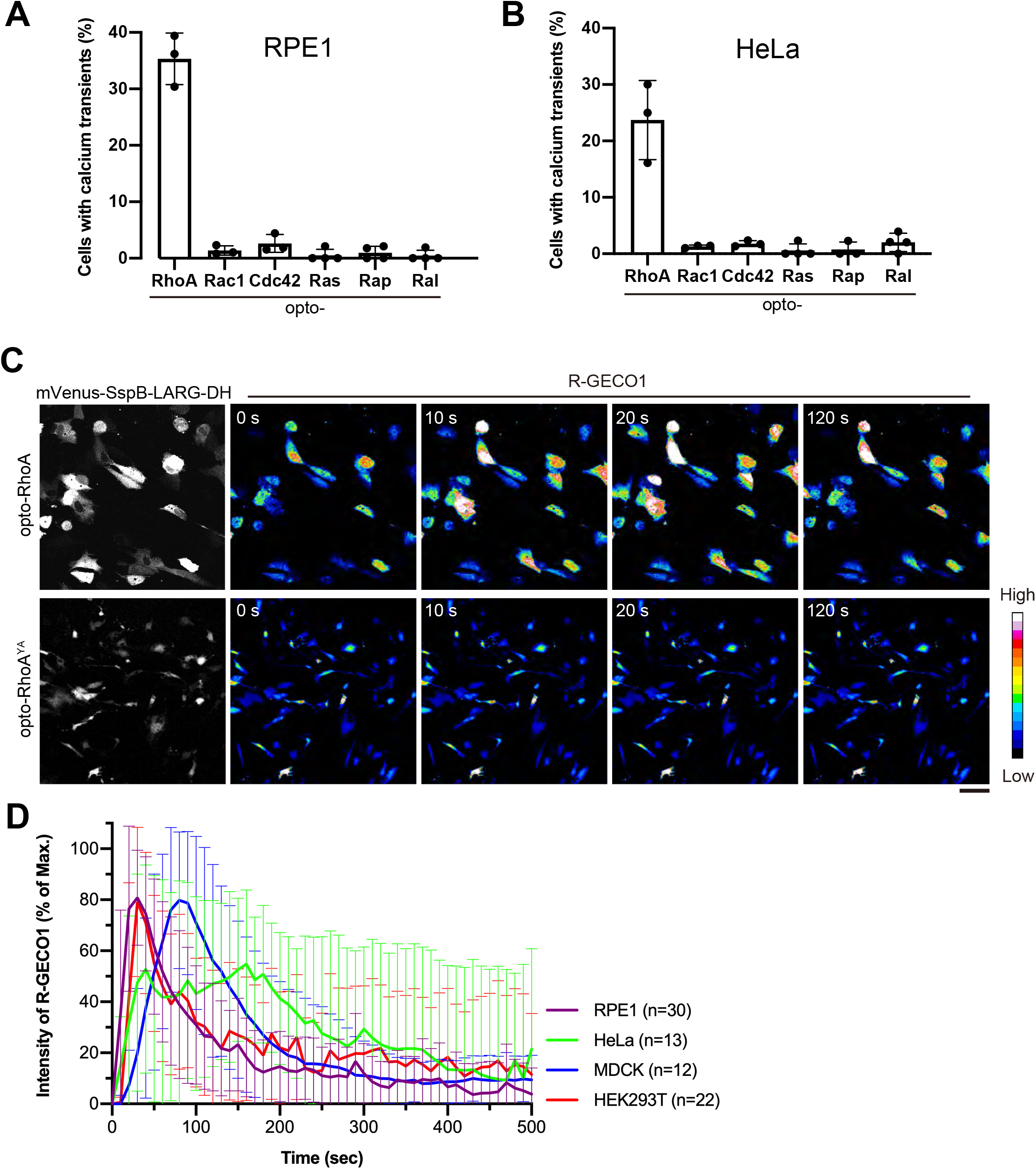
RhoA activation induces intracellular calcium transients in various cell types. **(A, B)** Screening for small GTPases that induce intracellular calcium mobilization. RPE1 **(A)** and HeLa **(B)** cells transiently express opto-GTPases, and the calcium reporter R-GECO1 were observed with confocal microscopy in Leibovitz’s L-15 medium. Opto-GTPases were activated by a multi-argon 458-nm laser every 10 s for 2 min. Percentages of cells exhibiting intracellular calcium transients (F/F_0_ > 2) in response to opto-GTPases activation are shown. Data are presented as means ± SD from more than three independent experiments. In total, >150 cells were analyzed for each photoswitch. **(C)** Representative images of RPE1 cells expressing opto-RhoA or GEF-inactive mutant opto-RhoA^YA^ and calcium reporter, R-GECO1. From time 0 s, opto-RhoA was activated by a multi-argon 458-nm laser every 10 s. Scale bar = 50 μm. **(D)** Intensity changes of R-GECO1 during RhoA activation in various cell types. Only cells exhibiting intracellular calcium transients were analyzed. The maximum fluorescence intensity of R-GECO1 in individual cells was set to 100%, and the intensity at 0 s was set to 0%. Data are presented the mean ± SD from indicated numbers of cells.

Further, we examined RhoA-mediated calcium transients in MDCK and HEK293T cells. These cells also exhibited the calcium transients immediately after RhoA activation (30 % and 41 % of cells in MDCK (n = 50) and HEK293T (n = 52) cells, respectively; **Supplementary Figure S5 and Movie 2**). Thus, all cells examined exhibited RhoA-mediated calcium transients. Of note, membrane blebbing, produced by actomyosin-mediated cellular contraction induced by RhoA activation (34), was seen in opto-RhoA-transfected HEK293T cells even before blue-light irradiation (**Supplementary Figure S5 and Movie 2**). This observation suggests that opto-RhoA was somewhat leaky; that is, opto-RhoA exerts background activity in the dark. The number of blebs was dramatically increased after the blue-light irradiation, indicating that the blue-light irradiation did increase RhoA activity.

Time course analysis showed that initiation of an increase in intracellular calcium concentration was observed within 10 s in RPE1, HeLa, and HEK293T cells; meanwhile, maximum concentration was observed about 20 s after light irradiation (**Figure 3D**). Among them, HeLa cells exhibited variability in the timing of calcium elevation. Meanwhile, high calcium concentrations were sustained longer in HeLa cells than those in other cell lines. In contrast, initiation of this increase was observed within 20–30 s, and the maximum increase was reached about 2 min after the light irradiation in MDCK cells (**Figure 3D**). Thus, cellular responses differed among cell types.

### Mechanisms of RhoA-induced calcium transients are different among cell types

We investigated molecular mechanisms of RhoA-induced intracellular calcium transients using small molecule inhibitors (see **Figure 8** for the summary). Several calcium channels are activated by RhoA and its downstream factors (35–37). We initially examined the RhoA–ROCK–myosin II axis, the major pathway of the RhoA signaling pathway. Both the ROCK inhibitor Y-276322 and myosin II inhibitor blebbistatin efficiently inhibited RhoA-induced calcium transients in MDCK and HEK293T cells, but surprisingly, this was not observed in RPE1 and HeLa cells (**Figure 4**). Actomyosin-mediated cellular contraction may activate mechanosensitive (MS) calcium channels such as Piezo1 and transient receptor potential (TRP) family channels in MDCK and HEK293T cells.

**Figure 4.**
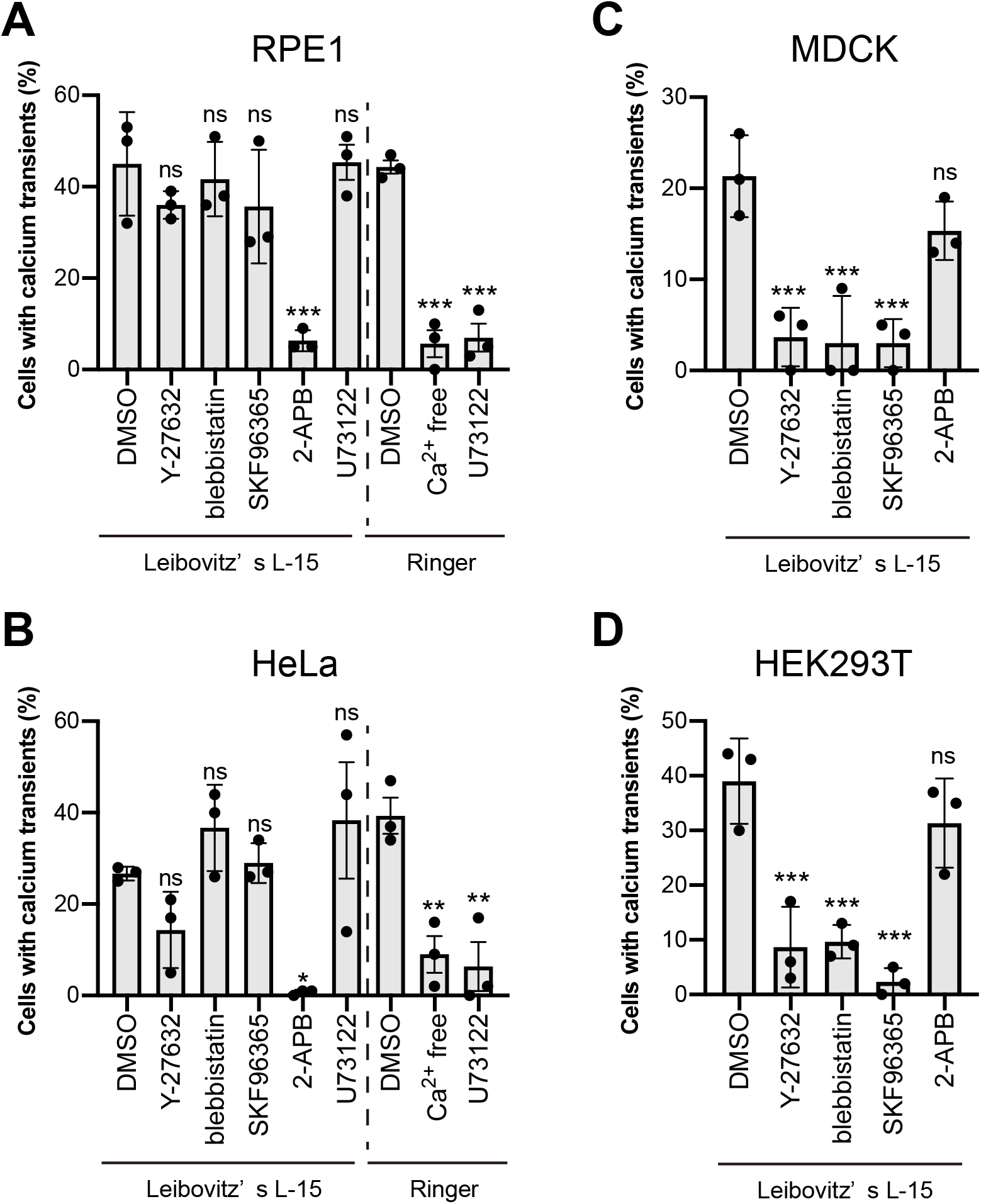
Mechanisms of intracellular calcium transients induced by RhoA are different among cell types. RPE1 (**A**), HeLa (**B**), MDCK (**C**), and HEK293T (**D**) cells expressing opto-RhoA and R-GECO1 were treated with DMSO, 50 μM Y-27632 (ROCK inhibitor), 20 μM blebbistatin (myosin II inhibitor), 10 μM SKF96365, 100 μM 2-APB (calcium channel inhibitors), or 10 μM U73122 (PLC inhibitor) and were observed using confocal microscopy (see Figure 3). Leibovitz’s L-15 medium and Ringer’s solution with or without Ca^2+^ were used for the experiments. Percentages of cells exhibiting intracellular calcium transients (F/F_0_ >2) in response to RhoA activation are shown. Data are shown as means ± SD from three independent experiments. In total, >200 cells were analyzed for each condition. Data were analyzed by one-way ANOVA followed by Dunnett’s test between DMSO-treated cells and other groups. ANOVA *F* = 9.473 and *p* = 0.0008 (**A**, Leibovitz’s L-15), *F* = 71.47 and *p* < 0.0001 (**A**, Ringer), *F* = 5.591 and *p* = 0.0069 (**B**, Leibovitz’s L-15), *F* = 16.65 and *p* = 0.0036 (**B**, Ringer), *F* = 14.62 and *p* = 0.0004 (**C**), *F* = 19.39 and *p* = 0.0001 (**D**). ***, *p* < 0.001; **, *p* < 0.01, *, *p* < 0.05; and ns, not significant; Dunnett’s test.

Further, we tested the non-selective calcium channel blockers SKF96365 and 2-APB (**Figure 4**). SKF96365 was observed to inhibit the RhoA-induced calcium transients in MDCK and HEK293T cells (**Figure 4C–D**). Conversely, 2-APB blocked transients in RPE1 and HeLa cells (**Figure 4A–B**). Thus, Mechanisms of RhoA-induced calcium transients are clearly different among cell types. 2-APB inhibits IP3R, and many small GTPases have been reported to activate PLC. However, as demonstrated in this study, only RhoA induced calcium transients among six small GTPases. We then focused on the molecular mechanisms of RhoA-induced calcium transients in RPE1 and HeLa cells to clarify this discrepancy.

### RhoA-PLCε axis induces calcium transients in RPE1 and HeLa cells

2-APB inhibits IP3R, and RhoA directly activates PLCε (35). Thus, we hypothesized that the RhoA–PLCε axis has induced calcium transients in RPE1 and HeLa cells. We also examined the effect of the PLC inhibitor U73122 **(Figure 4A–B)**. Initially, calcium transients were not affected by U73122 in L-15 medium. However, U73122 is reported to spontaneously form conjugates with the chemical components of cell culture medium, such as L-glutamine, glutathione, and bovine serum albumin. This conjugation causes a loss of inhibitory activity toward PLC (38). Calcium transients were abolished by U71322 in both RPE1 and HeLa cells when incubated in Ringer’s solution. As hypothesized, RhoA activates PLC, which in turn induces calcium transients.

We also measured RhoA-induced calcium transients in calcium-free buffers (**Figure 4A–B**). The percentage of cells exhibiting calcium transients significantly decreased in both cell types. However, about 10% of cells still showed transients, and intracellular calcium stores have been identified as a source of these calcium transients. Extracellular calcium could be required for proper calcium homeostasis.

We also knocked down PLCε in RPE1 and HeLa cells with small interference RNA (siRNA) and examined calcium transients (**Figure 5A–B**). RhoA activation by opto-RhoA in PLCε-depleted cells did not induce calcium transients. Further, ECFP-fused mouse PLCε (msPLCε), not ECFP alone, rescued calcium transient phenotypes in PLCε-depleted cells (**Figure 5C–E, Supplementary Figure S6**). PLCε is thus essential for calcium transients induced by RhoA in RPE1 and HeLa cells.

**Figure 5.**
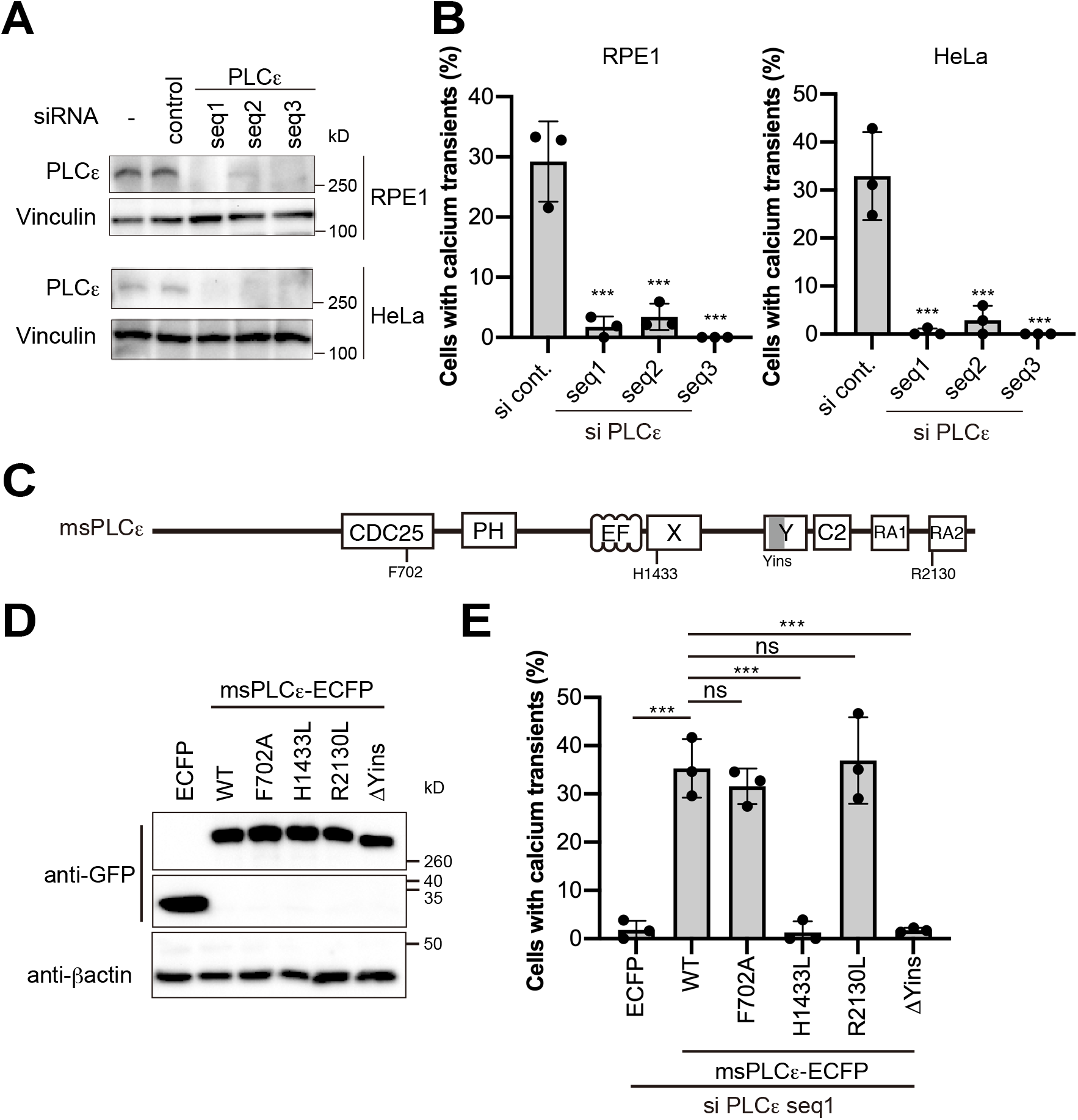
RhoA activates PLCε and induces intracellular calcium transients in RPE1 and HeLa cells. **(A)** RPE1 and HeLa cells were treated with PLCε-specific or control siRNAs for 48 h. Proteins in total cell extracts were separated with SDS-PAGE and immunoblotted with anti-PLCε and anti-vinculin antibodies. Vinculin was used as a loading control. **(B)** siRNA-mediated PLCε-depleted RPE1 and HeLa cells express opto-RhoA and R-GECO1 (see Figure 3). **(C)** Schematic of mouse PLCε (msPLCε). CDC25, CDC25 homology guanine nucleotide exchange factor (GEF) domain; PH, pleckstrin homology domain; EF, EF-hand motif; X and Y, X and Y domains fold to form the catalytic core of the phospholipase; C2, C2 domain; RA, ras (RA) association homology domain. **(D)** Proteins in total extracts from HEK293T cells expressing ECFP, msPLCε-ECFP wild-type (WT), or indicated msPLCε-ECFP mutants were separated with SDS-PAGE and immunoblotted with anti-GFP and anti-β-actin antibodies. β-actin was used as a loading control. The top and bottom images were from the same blot, but the middle image was obtained from another blot using the same sample because the molecular size was different. **(E)** siRNA-mediated PLCε-depleted HeLa cells transiently expressing ECFP, msPLCε-ECFP WT or indicated msPLCε-ECFP mutants, and R-GECO1 (see Figure 3). Data are presented as means ± SD from three independent experiments. In total, >150 cells were analyzed for each condition. Data were analyzed using one-way ANOVA followed by Dunnett’s test between control siRNA-treated cells (**B**) or msPLCε-ECFP WT-expressing cells (**E**) and other groups. ANOVA *F* = 44.28 and *p* < 0.0001 (**B**, RPE1), *F* = 32.70 and *p* < 0.0001 (**B**, HeLa), *F* = 42.33 and *p* < 0.0001 (**E**). ***, *p* < 0.001; and ns, not significant; Dunnett’s test.

We performed additional rescue experiments with several PLCε mutants in PLCε-depleted HeLa cells to further confirm RhoA–PLCε functions. PLCε features a lipase domain that includes a PH domain, four EF-hand domains, a catalytic core X-Y domain, and a C2 domain, as well as a CDC25 homology domain that activates Rap1 (39–41) (**Figure 5C**). This information led us to examine the essential function of PLCε for RhoA-induced calcium transients. The GEF-inactive mutant F702A PLCε, which corresponds to the F929A GEF-inactive mutant of RasGEF SOS (42), rescued calcium phenotype in PLCε-depleted cells. Conversely, the lipase-inactive mutant H1433L (39) did not rescue the phenotype (**Figure 5D–E**). Therefore, lipase activity appears essential for the generation of calcium transients.

PLCε is also regulated by Ras (43, 44), Rap1 (40), Ral (45), and heterotrimeric G-protein β and γ subunits (G_βγ_) (46). A unique insert in the Y domain (Yins in **Figure 5C**) of PLCε is required for its activation by RhoA (35). By contrast, Ras and Rap1 directly bind to the second Ras association domain (RA2 domain in **Figure 5C**) (47) of PLCε. Then, we performed rescue experiments in PLCε-depleted HeLa cells with a Yins-deleted PLCε mutant (ΔYins) and the R2130L PLCε mutant, which abolishes interactions with Ras (47) (**Figure 5D**). As a result, the R2130L mutant, but not the ΔYins mutant, rescued the calcium phenotype in PLCε-depleted cells (**Figure 5E**). These data suggest that RhoA activates the phospholipase activity of PLCε, which in turn induces intracellular calcium transients.

### RhoA activation at the plasma membrane but not at the Golgi is essential for the calcium transients

PLCε was recently reported to be activated at the Golgi, but not at the plasma membrane, in cardiac cells to hydrolyze phosphatidylinositol 4-phosphate [PI(4)P] (48–50). Our optogenetic construct iLID-CAAX^KRas^ largely localizes to the plasma membrane, but it also partially localizes to the Golgi. Thus, opto-RhoA could activate Golgi RhoA. However, PI(4)P hydrolysis produces inositol bisphosphate (IP_2_) but not IP_3_, and PLCε activation at the Golgi is unlikely to induce IP3R-dependent calcium release from the ER. We examined this issue by constructing opto-RhoA-TGN (*trans*-Golgi network) using the N-terminal transmembrane domain of 2,6-sialyltransferase (51) and opto-RhoA-Golgi carrying the KDELr D193N dominant-negative mutant (KDELr DN) (52) instead of the CAAX^KRas^ motif (**Figure 6A–B**). In both RPE1 and HeLa cells, only approximately 10% of cells expressing opto-RhoA-TGN exhibited calcium transients, whereas this was not observed in cells expressing opto-RhoA-Golgi (**Figure 6C**). Thus, RhoA activation at the plasma membrane appears essential for the generation of calcium transients.

**Figure 6.**
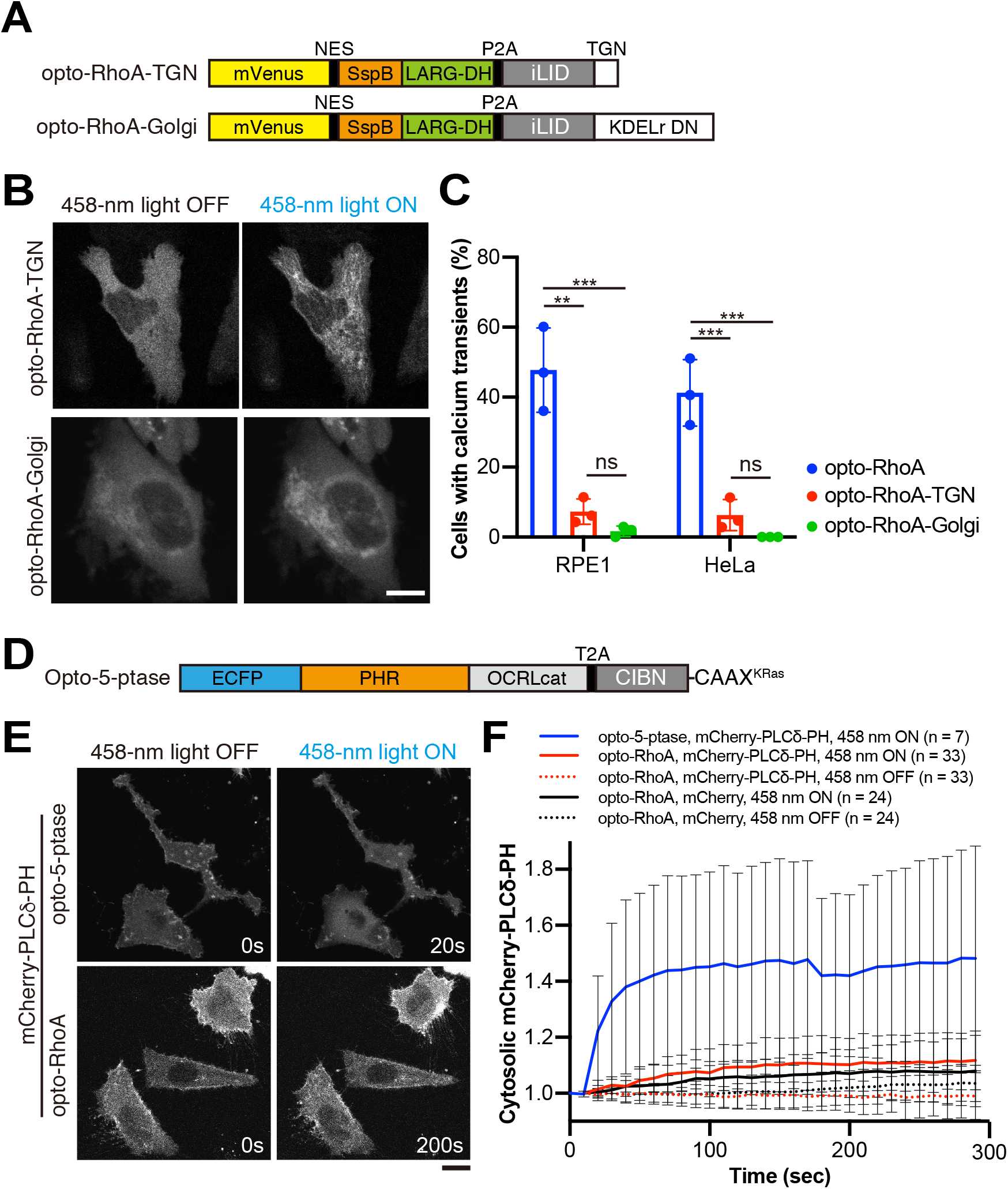
RhoA activates PLCε at the plasma membrane and marginally affects membrane phosphoinositide dynamics. **(A)** Schematic of opto-RhoA-TGN and opto-RhoA-Golgi. TGN, trans-Golgi network localized N-terminus transmembrane domain of 2,6-sialyltransferase; KDELr DN, dominant-negative (DN) KDELr D193N. **(B)** Representative images of HeLa cells expressing opto-RhoA-TGN or opto-RhoA-Golgi before (458-nm light OFF) and after (458-nm light ON) light irradiation with a 458-nm laser. Scale bar = 20 μm. **(C)** RPE1 and HeLa cells expressing opto-RhoA, opto-RhoA-TGN, or opto-RhoA-Golgi, and R-GECO1 (see Figure 3). Data are presented as means ± SD from three independent experiments (in total >150 cells were analyzed for each photoswitch). Data were analyzed using one-way ANOVA followed by Tukey’s test. ANOVA *F* = 35.41 and *p* = 0.0005 (RPE1), *F* = 40.38 and *p* = 0.0003 (HeLa). ***, *p* < 0.001; **, *p* < 0.01; and ns, not significant; Tukey’s test. **(D)** Schematic of opto-5-ptase. OCRLcat, inositol 5-phosphatase catalytic domain of OCRL. **(E)** Representative images of cells expressing opto-5-ptase or opto-RhoA and PI(4,5)P_2_ marker, mCherry-PLCδ-PH, before and after 458-nm light irradiation. Scale bar = 20 μm. **(F)** Quantification of cytosolic mCherry-PLCδ-PH intensity. HeLa cells expressing indicated photoswitches (opto-5-ptase or opto-RhoA), and markers (mCherry-PLCδ-PH or mCherry alone) were observed by confocal microscopy with or without 458-nm light irradiation every 10 s (458-nm ON or 458-nm OFF, respectively). Cells were first observed without 458-nm light irradiation, and then the same cells were observed with light irradiation. Data are presented the mean ± SD from the indicated number of experiments.

We next explored PI(4,5)P_2_ dynamics on the plasma membrane using the PI(4,5)P_2_ marker, a PH domain of PLCδ (PLCδ-PH) (53) and an optogenetic tool for controlling the inositol 5-phosphatase (opto-5-ptase) with the inositol 5-phosphatase domain of OCRL (OCRL_cat_) and the CRY2-CIBN heterodimerization system, as previously reported (54) (**Figure 6D**). This latter construct served as a positive control. CRY2-CIBN heterodimerization is induced by blue-light irradiation, and cytosolic ECFP-CRY2-OCRL_cat_ then translocates to the plasma membrane in functional form. When 5-phosphatase was recruited to the plasma membrane, the fluorescence intensity of cytosolic mCherry-PLCδ-PH increased to about 40% within 30 s (**Figure 6E–F and Movie 3)**, indicating that membrane PI(4,5)P_2_ decreased. In contrast, when RhoA was activated by opto-RhoA, the intensity of cytosolic mCherry-PLCδ-PH increased only about 10% even after 5 min. This result is comparable to intensity changes of mCherry alone (**Figure 6E–F and Movie 4**). RhoA-mediated cellular contraction might also cause this increase of mCherry intensity since we quantified mCherry intensity in a constant square area. Thus, the RhoA-PLCε axis functions at the plasma membrane but only marginally affects membrane phosphoinositide dynamics.

### RhoA–PLCε axis induces calcium signaling

Finally, we examined RhoA–PLCε-mediated intracellular calcium transients for activation of intracellular calcium signaling with an NFAT nuclear translocation assay. NFAT is identified as a transcription factor, and, after dephosphorylation by calmodulin-calcineurin, it translocates from the cytoplasm to the nucleus (55). Approximately 30% of cells exhibited mCherry-NFAT nuclear translocation after RhoA activation by opto-RhoA (**Figure 7A–B and Movie 5**). This response is comparable to the percentage of cells that exhibited calcium transients. Translocation was abolished by treatment with PLCε-specific siRNA (**Figure 7C–D**). Furthermore, RhoA-mediated NFAT translocation was abolished by pre-treatment with the calcium chelator, BAPTA-AM or treatment with calcineurin inhibitor cyclosporin A (**Figure 7E–F**). Thus, the RhoA–PLCε axis induces intracellular calcium signaling.

**Figure 7.**
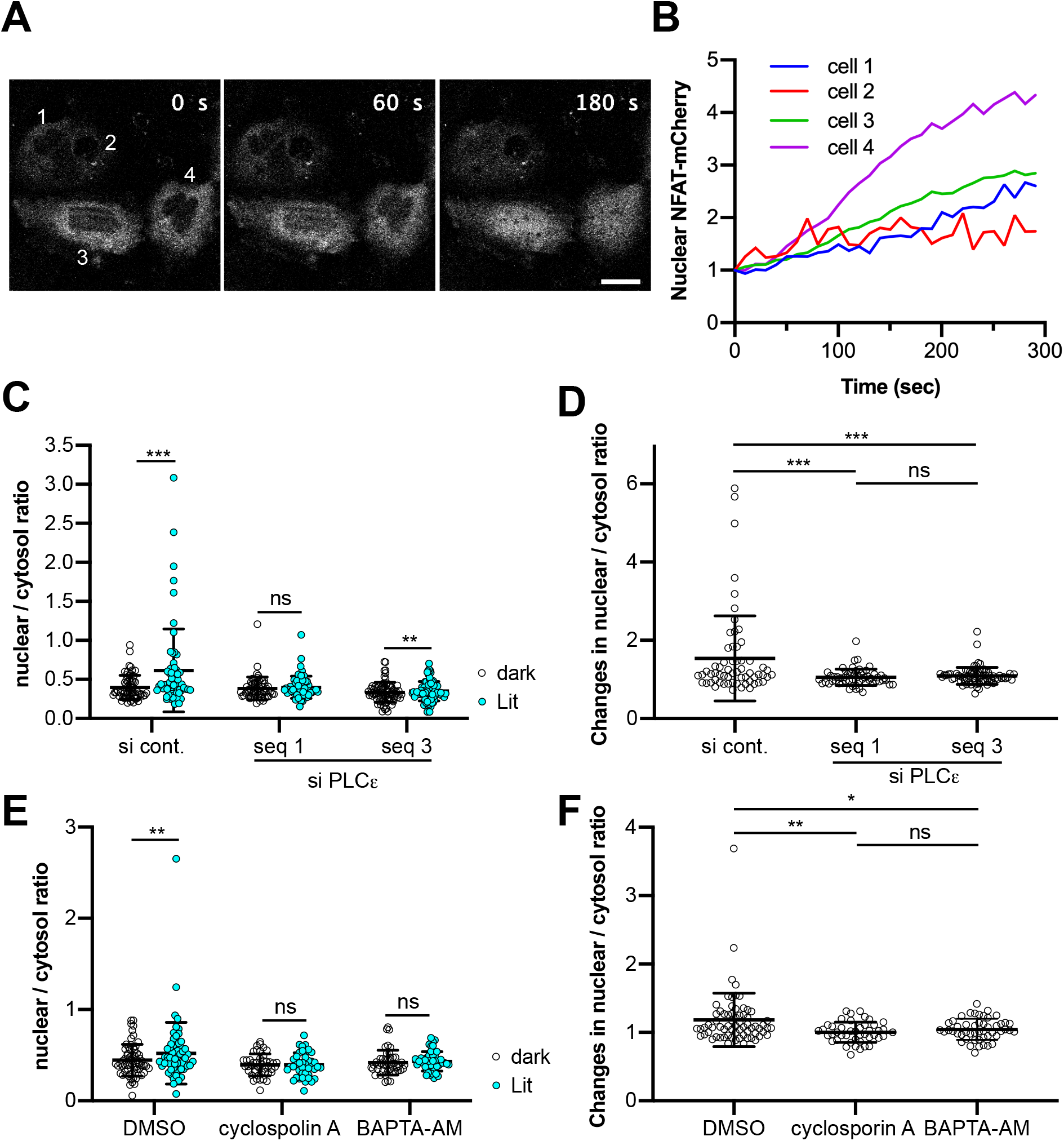
Optogenetic RhoA activation induces intracellular calcium signaling. **(A)** Time-lapse imaging of NFAT-mCherry translocation in HeLa cells. Cells expressing opto-RhoA and NFAT-mCherry were observed by confocal microscopy with a 458-nm laser irradiation every 10 s. Bar = 20 μm. **(B)** Time course of the relative fluorescence intensity of nuclear NFAT-mCherry. Cells indicated in **A** were analyzed. **(C–F)** Nuclear to cytosol ratio (**C, E**) and changes of this ratio (**D, F**) of NFAT-mCherry before (dark) and after 5 min (Lit). HeLa cells were treated with individual siRNA duplexes for 48 h and transiently expressed opto-RhoA and NFAT-mCherry (**C–D**). HeLa cells transiently expressing opto-RhoA and NFAT-mCherry were treated with DMSO, 10 μM cyclosporin A, or pre-treated with 2 μM BAPTA-AM for 1 h (**E–F**). N = 61, 58, and 72 cells (**C–D**), and N = 69, 44, and 50 cells (**E–F**). Data are presented as individual points and the mean ± SD from three independent experiments. Data were analyzed using a two-tailed paired Student’s *t*-test (**C, E**) or one-way ANOVA, followed by Tukey’s multiple comparisons test (**D, F**). ANOVA *F* = 11.23 and *p* < 0.0001 (**D**), *F* = 6.569 and *p* = 0.0018 (**F**). ***, *p* < 0.001; **, *p* < 0.01; *, *p* < 0.05 and ns, not significant; two-tailed paired Student’s t-test (**C, E**) and Tukey’s test (**D, F**).

## Discussion

In this study, we have constructed optogenetic tools using the iLID system to control Rho and Ras family GTPase activity (**Figures 1, 2**). We then performed cell-based systematic functional screening (**Figure 3**). Although we focused on the crosstalk between Rho/Ras family GTPases and the intracellular calcium signaling in this study, this approach for cell-based screening can be applied to other signaling pathways as well if both optogenetic tools and biosensors are available. Before introducing optogenetic tools, single cell-based enzymology was difficult due to the insufficient spatiotemporal resolution and specificity. Optogenetics has resolved this issue by exerting control over specific signaling molecules using a light stimulus with high spatiotemporal resolution. This tool provides a powerful platform for cell-based enzymology.

As intended, opto-GTPases specifically activated their target small GTPases (**Figure 2 and Supplementary Figure S3–S4**). In addition, opto-Cdc42 activated RhoA and Rac1, and opto-Ras activated RalB. As already discussed in the Results, these small GTPases are likely to be activated indirectly via Cdc42 and Ras activation. Note that these results do not mean that Rac1 and RhoA are activated in the same region of cells in which Cdc42 is activated. The crosstalk between Rho family small GTPases could be more complicated. It has been reported that when Cdc42 is locally activated using optogenetic tools, Rac1 was also activated on the same side of the cells, but the spatial distribution of Cdc42 and Rac1 activity differed (22, 31). Conversely, RhoA was activated on the opposite side of the cells (31). Thus, our results only illustrated that Rac1 and RhoA were activated via Cdc42 activation when measuring the entire cell.

The time course of small GTPase activation and inactivation by opto-GTPases differed among small GTPases (**Figure 2**). Conversely, with the exception of opto-Rap, the time course of the membrane translocation of opto-GTPases did not substantially differ (**Supplementary Figure S1–S2**). It is also known that upon EGF stimulation, Rho family small GTPases are rapidly activated, whereas Ras and Rap are gradually activated (56). Therefore, it is suggested that the ON–OFF kinetics of Rho family GTPases is rapid, and that of Ras family GTPases is slow in cells even with direct activation by their specific GEFs.

In our screening using opto-GTPases, only RhoA activation was seen to induce calcium transients (**Figure 3**). This was surprising because other GTPases are also reported to induce calcium signaling (11, 12). Regarding the reasons for this discrepancy, the sensitivity of our experiments may not have been sufficient to detect calcium transients by such GTPase activation, or leaks in our optogenetic systems (**Supplementary Figure S5**) may have affected signaling responses. However, RhoA activation has induced calcium transients under the same conditions. Another possibility is that activation of only individual small GTPases does not induce calcium transients. Other signals, such as PI3K, could be required for activating calcium signaling. In any case, our study was the first to induce temporal activation of specific GTPases and monitor subsequent intracellular calcium changes. This approach is expected to have much higher specificity and time resolution compared with previous studies. Further, our systems can simultaneously stimulate other signals using multiple optogenetic tools, and we can examine the dual activation of small GTPases and other signals for the induction of calcium transients. Such experiments could explain many discrepancies in signal transduction.

RhoA-induced calcium transients have been reported in only a few cell types (35–37), and therefore, they have not received substantial attention. Because RhoA-induced calcium transients were also observed in all cell lines examined in this study (**Figure 3 and Supplementary Figure S5**), which has not been studied before, RhoA appears to induce calcium signaling generally.

Unexpectedly, the molecular mechanisms underlying the generation of calcium transients were found to notably differ among cell types (**Figures 4 and 8**). In MDCK and HEK293 cells, the RhoA–ROCK–myosin II axis induced calcium transients (**Figure 4C–D**). Actomyosin-mediated contractile force is also reported to induce membrane tension changes that activate MS channels (57) (**Figure 8, lower scheme**). One candidate is the Piezo channel reported in both MDCK (58) and HEK293T cells (59). However, Piezo channels are not sensitive to SKF96365 (60), which inhibited transient induction (**Figure 4C–D**). Thus, other MS calcium channels, such as a member of the TRP family of channels, may be needed for RhoA-induced calcium transients. Further studies are also required to elucidate molecular mechanisms of RhoA–ROCK–myosin II-mediated calcium transients.

**Figure 8.**
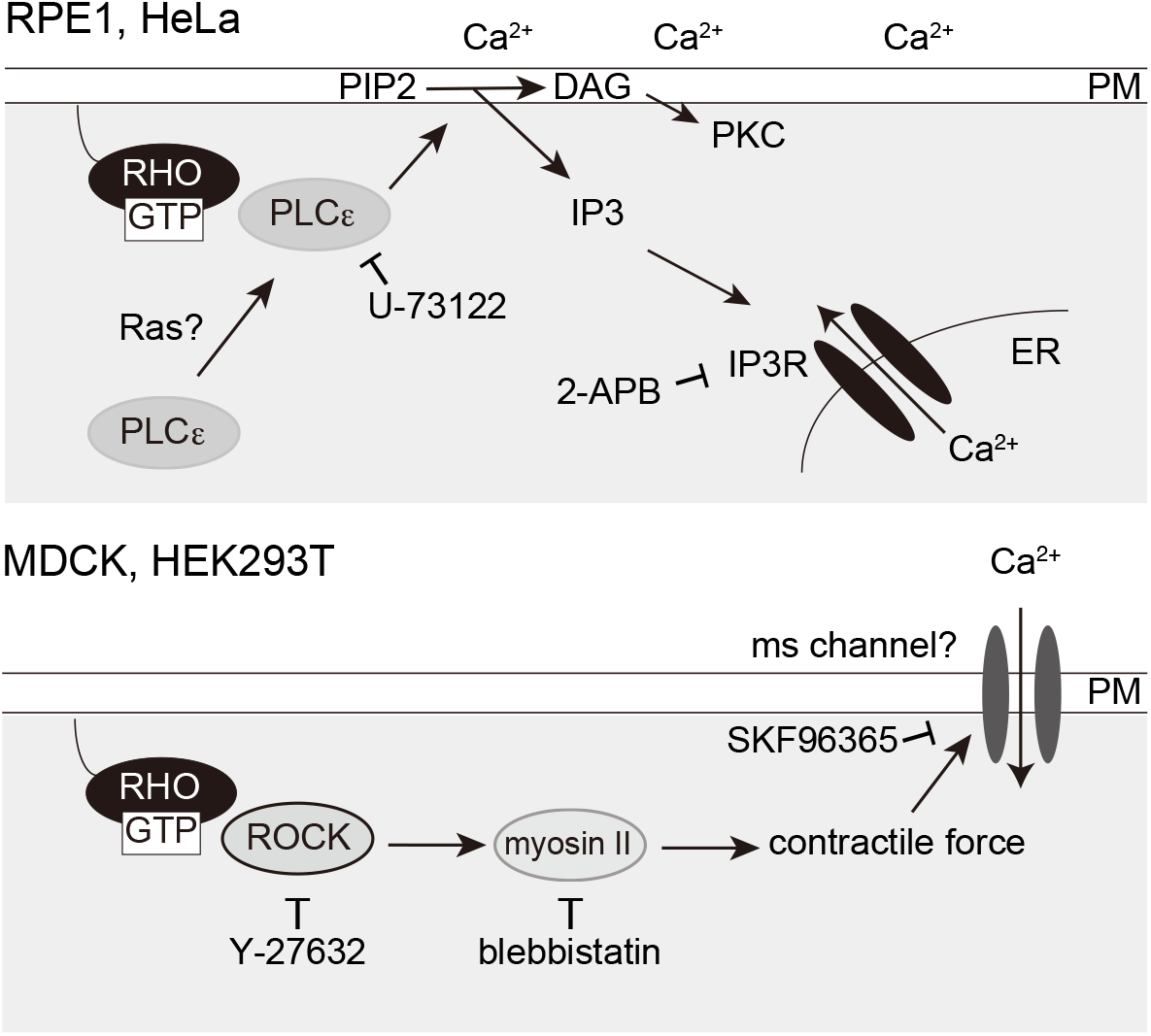
Summary of the study. RhoA activation induces calcium transients in all examined cell types. However, the molecular mechanisms differed among the cell types. In RPE1 and HeLa cells, the RhoA–PLCε–IP3R axis induced Ca^2+^ release from the ER (**Top**). Because PLCε also produces diacylglycerol (DAG), this pathway can also activate PKC signaling. In addition, extracellular calcium was required for this process. Ras did not induce calcium transients, but it may be required for PLCε recruitment to the plasma membrane. Conversely, the RhoA–ROCK–myosin II axis induced calcium transients (**Bottom**). Actomyosin may exert a contractile force that which activates a mechanosensitive (MS) channel. Small inhibitors (U-73122, 2-APB, Y-27632, blebbistatin, and SKF96365) that blocked calcium transients are presented next to the target molecules. PM, plasma membrane.

By contrast, the RhoA–PLCε axis was determined to be functional in RPE1 and HeLa cells (**Figures 4, 5 and 8, upper scheme**). Calcium transients were totally blocked by 2-APB and partially observed in Ca^2+^-free buffer, and transients were apparently induced by the IP_3_-IP3R pathway that promotes Ca^2+^ release from the ER. Nevertheless, the percentage of cells exhibiting transients has decreased in Ca^2+^-free butter compared with cells in Ca^2+^-containing buffer. This finding may reflect changes in cellular responsiveness since cell processes, such as cell-substrate adhesion, membrane lipid composition, and intracellular calcium homeostasis, are largely dependent on extracellular Ca^2+^, for example, purified PLCε exhibits Ca^2+^-dependent phospholipase activity (44). It is also possible that calcium influx from the extracellular space is essential for this process.

PLCε has been identified to be activated by other small GTPases, including Ras, Rap, and Ral (40, 44, 45), which is inconsistent with the present finding that these GTPases did not induce calcium transients (**Figure 3**). This discrepancy can be explained by the fact that there is no evidence that small GTPases directly activate the phospholipase activity of PLCε. In previous studies, phospholipase activity was typically examined as IP_3_ accumulation in COS-7 cells co-expressing constitutively active mutants of small GTPases and PLCε exogenously (35, 44, 45, 61–63). This assay cannot exclude the side effects of downstream signaling. In addition, the purified KRas G12V constitutively active mutant the activated phospholipase activity of N-terminally deleted PLCε (64), but it did not stimulate the activity of the full-length intact PLCε (43) in an *in vitro* reconstitution assay. However, there is much evidence that active Ras and Rap interact with the RA2 domain of PLCε and induce translocation from the cytoplasm to the plasma membrane and Golgi, respectively (40, 43). Bunney et al have been proposed a two-step activation model for PLCε in which Ras both recruits PLCε to the plasma membrane and partly releases the autoinhibition by the RA2 domain, and another molecule such as RhoA is required for full PLCε activation (47). Our data could support this model because RhoA is more likely to be decisive for activating the phospholipase activity of PLCε rather than Ras, Rap, and Ral. Although only RhoA activation was sufficient to activate PLCε in our experiments, whether RhoA alone is actually sufficient for PLCε activation must be carefully considered. Because it has been reported that RhoA could not activate RA2 domain-depleted PLCε. However, it can be activated by RhoA via fusing the C-terminus of the CAAX motif; RA2 domain-depleted PLCε-CAAX then localizes to the plasma membrane via CAAX (61). Therefore, we attempted to determine whether RhoA also recruits PLCε to the plasma membrane, but definitive results were not obtained. It may be possible that the activity of endogenous Ras or other factors was sufficient for the recruitment, and therefore, RhoA activation alone activated PLCε in our experimental conditions. Thus, further studies are needed to understand the molecular mechanisms of PLCε activation by RhoA, including whether the translocation of PLCε mediated by Ras or Rap1 is essential.

We revealed that RhoA-mediated calcium transients induce NFAT translocation to the nucleus (**Figure 7**), indicating that RhoA-induced calcium transients activate intracellular calcium signaling. Because both RhoA and calcium signaling commonly regulate actin cytoskeleton reorganization, cell proliferation, and gene expression (3–5, 65, 66), signaling coordination appears reasonable. One of the possible physiological roles of RhoA-mediated calcium transients is related to actomyosin-mediated cellular contraction. Both Rho/ROCK and Ca^2+^ activate MLC. ROCK phosphorylates and inactivates MLC phosphatase and also directly phosphorylates MLC, whereas Ca^2+^/calmodulin phosphorylates and activates MLC kinase (6, 7). For example, vascular smooth muscle contraction and endothelial cell contraction, which are required for endothelial permeability, are induced by thrombin (67, 68). Thrombin is known to activate Gα_12/13_, which in turn activates RhoA. It has been reported that both Ca^2+^ and the RhoA/ROCK pathway are necessary for the full thrombin response (68), and the present study suggests that these signaling pathways could be integrated into RhoA. As another example, actomyosin-mediated cellular contraction has essential roles during cytokinesis (69). It has been reported that calcium transients were observed at the equatorial region during cytokinesis and were required for efficient ingression of the cleavage furrows (70, 71). Ca^2+^ release from the ER via IP3R has been suggested as the Ca^2+^ source during cytokinesis, but the molecular mechanisms have remained unclear (70). It is possible that RhoA induces calcium transients at the equatorial region during cytokinesis because RhoA is a master regulator of cleavage furrow formation and ingression (72).

In addition to thrombin, thromboxane A2, LPA, and sphingosine-1-phosphate (S1P) are G protein-coupled receptors (GPCR) ligands that have been well established as RhoA activators. The GPCR–Gα_12/13_–RhoA axis both regulates cytoskeletal rearrangement and activates transcription factors such as activator protein-1 (AP-1), NFκB, myocardin-related transcription factor A (MRTF-A), and Yes-associated protein (YAP), as well as regulates cell proliferation, differentiation, and inflammation (73). Because calcium signaling also regulates various transcription factors, the combinational regulation of Rho- and calcium-dependent gene expression is possible. Indeed, AP-1 and NFAT form a complex and induce the expression of genes required for immune responses such as IL-2 (74). Because the downstream signals and their physiological roles would vastly differ among cell types, the physiological functions of RhoA-induced calcium transients should be examined in each cell type in the future.

## Experimental procedures

### Antibodies

Commercial antibodies and their dilution were indicated as follows; mouse anti-GFP antibody diluted at 1:1000 (clone mFX75, FUJIFILM Wako Pure Chemical Corp., Osaka, Japan); anti-vinculin antibody diluted at 1:1000 (clone 2B5A7, Proteintech Group, Chicago, IL, USA); rabbit anti-PLCE1 antibody diluted at 1:1000 (HPA015597, Atlas Antibodies, Bromma, Sweden); peroxidase-conjugated mouse anti-β-actin antibody diluted at 1:10,000 (clone 2F3, FUJIFILM Wako Pure Chemical Corp.); peroxidase-conjugated sheep anti-mouse IgG antibody diluted in 1:4000 (NA931, GE Healthcare, Little Chalfont, UK); and peroxidase-conjugated donkey anti-rabbit IgG antibody diluted in 1:2000–1:4000 (NA934, GE Healthcare).

### Cell culture and transfection

hTERT-immortalized human retinal pigment epithelial (RPE1) cells (ATCC CRL-400) were cultured in DMEM and F12 nutrient mix (1:1), supplemented with 10% (v/v) fetal bovine serum (FBS: Biowest, Nuaillé, France). HeLa (RCB0007: RIKEN BRC through the National BioResource Project of the MEXT/AMED, Japan), MDCK (kindly gifted from Dr. N. Yui), and HEK293T (obtained from TaKaRa, Shiga, Japan) cells were cultured in DMEM supplemented with 10% FBS. Plasmid transfection was performed using Lipofectamine™ 2000 (Thermo Fisher Scientific, Waltham, MA, USA) or Viafect (Promega, Madison, WI, USA) following the manufacturer’s instructions. Reagents were obtained from commercial sources as follows: Y-27632 dihydrochloride (Focus, Biomolecules, Plymouth Meeting, PA, USA); (−)-blebbistatin; U73122 (Cayman Chemical, Ann Arbor, MI, USA); 2-APB; SKF96365 (FUJIFILM Wako Pure Chemicals); BAPTA-AM; and cyclosporin A (Nacalai Tesque, Kyoto, Japan).

### DNA construction

pLL7.0: Venus-iLID-CAAX (from KRas4B) and pLL7.0: tgRFPt-SSPB wild-type (WT) (19) were gifts from Brian Kuhlman (Addgene plasmid #60411 and #60415). PR_GEF (2XPDZ-mCherry-Larg (DH)) was provided by Michael Glotzer (Addgene plasmid #80407). CMV-R-GECO1 (32) and pDDRFP-A1B1-DEVD (75) were gifted by Robert Campbell (Addgene plasmid #32444 and #36294). pcDNA3-HA-human OCRL (76) was supplied by Pietro De Camilli (Addgene plasmid #22207). GFP-C1-PLCdelta-PH (53) was obtained from Tobias Meyer (Addgene plasmid #21179). pTriEx-RhoA FLARE.sc Biosensor WT (77) was gifted by Klaus Hahn (Addgene plasmid #12150). ERK2-GFP was gifted by Kai Zhang (30). Human C3G cDNA was procured from Hiroki Tanaka (Kyoto University). HA-NFAT1(1-460)-mCherry was constructed previously (78). Total RNA was obtained from RPE1 and HeLa cells using QIAzol Lysis Reagent (QIAGEN, Hilden, Germany) following the manufacturer’s protocol, and first-strand cDNA was synthesized with 1 μg of total RNA using a SuperScript III First-Strand Synthesis System for RT-PCR (TaKaRa) following the manufacturer’s protocol.

msPLCε cDNA was amplified by PCR from a mouse liver first-strand cDNA library (kindly gifted from Tomohiro Ishii) using Tks Gflex polymerase (TaKaRa) and was cloned between the *Nhe*I and *Kpn*I sites of pECFP-N1 (Clontech, Palo Alto, CA, USA). msPLCε sequences were sensitive to siPLCε seq1, and a siRNA-resistant msPLCε mutant was used in this study. All point mutations were generated using site-directed mutagenesis methods by Tks Gflex polymerase. msPLCε ΔYins mutant was generated with overlap extension PCR.

To create pmCherry-PLCδ-PH, EGFP was removed from pEGFP-C1 PLCdelta-PH and replaced with mCherry from pmCherry-C1 (TaKaRa). To create pmCherry-C1 RBD_rhotekin_, RBD_rhotekin_ was amplified from pTriEx-RhoA FLARE.sc Biosensor WT and cloned between the *Bgl*II and *Eco*RI site of pmCherry-C1. To create ERK2-mCherry, ERK2 cDNA from ERK2-EGFP was cloned between the *Xho*I and *Bam*HI site of pmCherry-N1 (TaKaRa).

Then, the DNA coding sequence was cloned between *Nhe*1 and *Not*1 of pEGFP-N1 to create opto-RhoA (mVenus-NES-SspB-LARG-DH-P2A-iLIDcaa x). The NES sequence (FGIDLSGLTL) was directly linked to mVenus. SspB and LARG-DH (DH domain of LARG, amino acids 766–986) were linked by LDSAGGSAGGSAGGLE. The P2A peptide sequence (GSG)ATNFSLLKQAGDVEENPGP was directly linked to LARG-DH and followed by the linker APGS to localize the 2A product protein to the precise subcellular domain (79).

The transmembrane domain of 2,6-sialyltransferase (amino acids 1–32) was then amplified from the HeLa cDNA library to create opto-RhoA-TGN, and KDELr was amplified from the RPE1 cDNA library by PCR to create opto-RhoA-Golgi. These fragments were connected to iLID by overlap extension PCR with a GSGSGS (3×GS) linker. iLID-KDELr was first cloned into a pECFP-C1 vector (Clontech), and KDELr D193N dominant-negative mutant (KDELr DN) was then generated by site-directed PCR. iLID-CAAX^Kras^ of opto-RhoA was replaced with iLID-TGN or iLID-KDELr DN.

The LARG-DH of opto-RhoA was replaced with a short linker (VDEFELDI), a DH-PH domain of Tiam (Tiam-DH-PH, amino acids 1012–1591), a DH-PH domain of ITSN (ITSN-DH-PH, amino acids 1230–1580), a catalytic domain of SOS2 (SOS2cat, amino acids 563–1048), a catalytic domain of C3G (C3Gcat, amino acids 687–1077), and a GEF domain of RGL2 (RGL2-GEF, amino acids 1–518) to create opto-control, -Rac1, -Cdc42, -Ras, -Rap, and -Ral, respectively.

Opto-5-ptase was constructed as previously reported for mCherry-PHR-iSH-2A-CIBNcaax (79), in which the vector backbone was changed to pEGFP-N1, and mCherry and iSH were replaced with ECFP and the inositol 5-phosphate domain of OCRL (OCRLcat, amino acids 234–539), respectively.

ddRFP-based small GTPase biosensors were constructed as described previously (29). To generate pRA-C1 and pB3-C1, ddRFPA1 and ddRFPB1 were amplified from pDDRFP-A1B1-DEVD and cloned between the *Nhe*I and *Bsr*GI or *Nhe*I and *Bsp*EI sites of the pEGFP-C1 vector, respectively. To create pRA-RhoA, pRA-Rac1, pRA-Cdc42, pRA-HRas, pRA-Rap1A and pRA-RalB, RhoA, Rac1, Cdc42, HRas, Rap1A, and RalB cDNAs, respectively, were cloned between the *Bsr*GI and *Bam*HI sites of pRA-C1. Then, RBD_rhotekin_-P2A-RA-RhoA, RBD_PAK1_ (amino acids 69–109)-P2A-RA-Rac1, RBD_PAK1_-P2A-RA-Cdc42, RBD_Raf1_ (amino acids 51–131)-P2A-RA-HRas, RBD_RalGDS_ (amino acids 788–885)-P2A-RA-Rap1A, and RBD_Sec5_ (amino acids 5–97)-P2A-RA-RalB were amplified by overlap extension PCR and cloned between the *Eco*RI and *Bam*HI sites of pB3-C1 to create pB3-RBD_rhotekin_-P2A-RA-RhoA, pB3-RBD_PAK1_-P2A-RA-Rac1, and pB3-RBD_PAK1_-P2A-RA-Cdc42, or the *Xho*I and *Bam*HI sites of pB3-C1 to create pB3-RBD_Raf1_-P2A-RA-HRas, pB3-RBD_RalGDS_-P2A-RA-Rap1A, and pB3-RBD_Sec5_-P2A-RA-RalB, respectively.

All cloned fragments were verified by sequencing.

### Small interference RNAs

Transfection of siRNA duplexes used Lipofectamine™ RNAiMAX reagent following the manufacturer’s protocol (Thermo Fisher Scientific). Each duplex was used at a final concentration of 10 nM. Double-strand RNAs were purchased from Ambion, Thermo Fisher Scientific. Target nucleotide sequences were 5′–GCAAGGAGCTGATCGATCT–3′ (s27658, siPLCε seq1), 5′–GGACATAGGCTGACAACCA–3′ (s27659, siPLCε seq2), and 5′–TACTGCGATATTGAAGTCC–3′ (s27660, siPLCε seq3), respectively. Negative control #2 siRNA (Silencer Select, Thermo Fisher Scientific) was used as a negative control.

### Live cell imaging

When only the plasmid transfection was performed, cells were seeded on glass-bottomed dishes (Greiner Bio-one, Kremsmünster, Austria) coated with collagen (Cellmatrix Type IC; Nitta Gelatin Inc., Osaka, Japan) 2 days before the observations. On the next day, cells were transfected with the plasmid and were analyzed a day later. When both siRNA and plasmid transfection were performed, cells were seeded 3 days before the observations. On the next day, siRNA transfection was performed. After 24 h since siRNA transfection, plasmid transfection was performed, and cells were analyzed a day later. Before experiments, the medium was changed to Leibovitz’s L-15 (Thermo Fisher Scientific), Ringer’s solution (138 mM NaCl, 5.6 mM KCl, 2mM CaCl_2_, 2 mM MgCl_2_, 4 mM D-glucose, 5 mM HEPES, and 2 mM sodium pyruvate (pH 7.4, adjusted with NaOH)), or calcium-free Ringer’s solution (sans CaCl_2_). Cells were analyzed under serum-starved conditions, and a closed heated chamber was used during live cell imaging at 37 °C without CO_2_. Images were obtained with a confocal laser scanning microscope (FV1200, Olympus, Tokyo, Japan) on an IX83 microscope (Olympus) equipped with 40×/0.95 NA and 20×/0.7 NA dry objective lenses and FV10-ASW software (Olympus). Excluding the images in Figure S1, photo-activation and imaging were performed with standard ECFP(C-Y-R), EYFP(C-Y-R), and DsRed2(C-Y-R) settings (Argon laser power, 10% at 458-nm and 1% at 515-nm; diode laser power, 1–10% at 559-nm; DM 458/515/560 dichroic excitation, SDM510 and SDM560 emission filters; 475–500 nm emission window for ECFP, 530–540 nm emission window for EYFP, and 570–670 nm emission window for DsRed2) using the sequential line capturing mode (scan rate: 10 μs per pixel [pixel size: 0.31 μm in Figures 1, 2, 6B, 6E, 6F, 7, and S2–S4; 1.24 μm in Figures 3, 4, 5, 6C, and S5–S6]). In Supplementary Figures S1A and C, photo-activation was performed with an Argon laser power of 1–10% at 458-nm or 1–10% at 515-nm, and the other settings were as previously mentioned. In Figure S1B, images and photo-activation were performed with standard EGFP and DsRed2 settings (1–10% 488-nm Argon laser power; 2% 559-nm diode laser power; DM 405/488/559/635 dichroic excitation, a SDM560 emission filter; 500–545 nm emission window for EGFP, and 570–670 nm emission window for DsRed2) using the sequential line capturing mode (scan rate: 10 μs per pixel [pixel size: 0.31 μm]). The time interval for both imaging and opto-activation was 20 s in Figure 1C–D and 10 s in all other figures. The mVenus images were obtained with the EYFP channel, and the mCherry, R-GECO1, and ddRFP images were obtained with the DsRed2 channel. Local irradiation in Figure 1C–D was performed using multi-area time-lapse software (Olympus).

### Image analyses

Image analyses were performed using ImageJ/Fiji software (NIH) (80).

The membrane translocation of opto-GTPases (Figure S1–S2) and membrane localization of mCherry-PLCδ-PH (Figure 6F) were evaluated according to changes in cytoplasmic fluorescence (79). The fluorescence signal intensity was measured in a 6.20 μm × 6.20 μm region in the cytoplasm throughout imaging. Then, the background fluorescence intensity was subtracted, and the relative fluorescence level was calculated by dividing the fluorescence intensity by that at 0 s (Figure 6F) or by that at 50 s (Figure S1–S2).

The following procedures were used to measure changes in the fluorescence intensity of ddRFP (Figures 2 and S3). First, binary images were produced, and cell masks were generated using the “Analyze Particles” command in ImageJ/Fiji. Next, the value was set to 1 for the inside of cells and NaN for the outside of cells and multiplied by the value of the original image, and fluorescence intensity was measured in the entire cell. Then, the background fluorescence intensity was subtracted, and the relative fluorescence level was calculated by dividing the fluorescence intensity by that at time 50 s.

The changes in R-GECO1 fluorescence intensity were measured in a manually decided constant square region in which the cells of interest entered as much as possible without overlapping other cells. Then, the background fluorescence intensity was subtracted, and the relative fluorescence level was calculated by dividing the fluorescence intensity by that at time 0 s. Cells with more than 2-fold increases in R-GECO fluorescence intensity (F/F_0_ > 2) were counted as cells with a calcium transient.

NFAT-mCherry translocation was evaluated according to changes in cytoplasmic and nuclear fluorescence. The fluorescence signal intensity was measured in a manually decided constant square region in the cytoplasm and nucleus throughout imaging. Then, the background fluorescence intensity was subtracted. The nuclear/cytosol ratio was calculated by dividing the nuclear fluorescence intensity by the cytosol fluorescence intensity at 0 and 5 min. The changes of the nuclear/cytosol ratio were calculated by dividing the nuclear/cytosol ratio at 5 min by that at 0 min.

ERK-mCherry translocation was evaluated on the basis of the changes in nuclear fluorescence, similar to NFAT-mCherry translocation described above.

Adobe Illustrator CS6 (Adobe Systems, San Jose, CA) was used for final figure preparation.

### Immunoblotting

Cells were lysed with 2 × SDS sample buffer (125 mM Tris-HCl (pH6.8), 4 % (w/v) SDS, 10 % (w/v) sucrose, 0.01 % (w/v) BPB, 5 % (v/v) 2-mercaptoethanol). Proteins were separated with SDS-PAGE and then later transferred to PVDF membranes by electrophoresis using a Bio-Rad Trans-Blot device (Bio-Rad, Richmond, CA). Blots were blocked for 1 h at room temperature or overnight at 4 °C with 5 % skim milk in Tris-buffer saline with Tween-20 (TBS-T, 50 mM Tris-HCl (pH 7.4), 138 mM NaCl, 2.7 mM KCl, and 0.1 % (v/v) Tween-20). Membranes were washed with TBS-T and incubated with indicated antibodies diluted in Can Get Signal Solution 1 (TOYOBO, Osaka, Japan) for 1 h at room temperature. Membranes were washed in TBS-T, incubated with appropriate horseradish peroxidase-conjugated secondary antibodies diluted in Can Get Signal Solution 2 (TOYOBO) for 1 h at room temperature, and washed again in TBS-T. Blots were developed with ImmunoStar Zeta (FUJIFILM Wako Pure Chemical), and chemiluminescence signals were visualized via a ChemiDoc MP system (Bio-Rad).

### Statistics

Statistical analyses were performed using Student’s *t*-test for comparisons between two samples (Figures 7C, 7E, and S5), and one-way ANOVA followed by Dunnett’s test (Figures 2C–E, 2G–H 4, 5B, 5E, and S4) or Tukey’s test (Figures 6B, 7D, and 7F) and Brown-Forsythe ANOVA test followed by Dunnett’s T3 (Figure 2F) for multiple comparisons, using Excel (Microsoft) and GraphPad Prism 9.0.0 (GraphPad Software, San Diego, CA, USA). *P* values < 0.05 were considered statistically significant. Graphs were drawn using GraphPad Prism 9.

## Data availability

All data are contained within this article and in the supporting information.

## Acknowledgments

The authors are grateful to Dr. Mari Ishigami-Yuasa (Chemical Biology Screening Center, TMDU) for technical helps, Dr. Naofumi Yui (TMDU) for the kind gift of MDCK cells, Dr. Toshiyuki Kakumoto for plasmid construction, the Research Core Center (TMDU) for usage of the ChemiDoc MP system, and Enago (www.enago.jp) for the English language review. We also thank Drs. Tomohiro Ishii and Toshifumi Asano for the helpful discussion and Satoko Nakamura for secretarial expertise.

## Abbreviations

DAG: diacylglycerol
ER: endoplasmic reticulum
GAP: GTPase activating protein
GEF: guanine nucleotide exchange factor
iLID: improved light-inducible dimer system
IP_3_: inositol-triphosphate
IP3R: IP3 receptor
LOV2: light-oxygen-voltage-sensing domain 2
MS: mechanosensitive
PLC: phospholipase C
PI(4)P: phosphatidylinositol 4-phosphate
PI(4,5)P_2_: phosphatidylinositol 4,5-bisphosphate
RA domain: Ras association domain
RBD: Ras/Rho binding domain
TGN: *trans*-Golgi network
TRP channel: transient receptor potential channel

