## Supplemental Figure legends for "Optogenetic control of small GTPases reveals RhoA-mediated intracellular calcium signaling"

Supplementary information for “**Optogenetic control of Small GTPases reveals RhoA-mediated intracellular calcium signaling**” by H. Inaba, Q. Miao and T. Nakata.

**Supplementary Figure Legends**

**Figure S1. Time courses of mCherry-SspB-LARG-DH translocation in response to irradiation with different wavelengths and laser power.** HeLa cells expressing the mCherry version of opto-RhoA were observed via confocal microscopy in Leibovitz’s L-15 medium. Cells were irradiated by a 458- (**A, D**), 488- (**B**), or 515-nm (**C**) laser at the indicated power every 10 s over the period of 60–170 s. Cytosolic mCherry-SspB-LARG-DH levels were quantified, and the fluorescence intensity was normalized by the intensity at 50 s. Data are presented as the mean ± SD. The same cells were observed in each panel. N = 5, 4, 5, and 5 cells, respectively.

**Figure S2. Time course of opto-GTPase translocation.** HeLa cells expressing the mCherry version of opto-GTPases were observed (Figure S1). Data are presented as the mean ± SD. N = 6, 5, 6, 7, 3, and 5 cells, respectively. A scale-modified and merged graph, including opto-RhoA (Figure S1), is presented to compare the kinetics of opto-GTPases (**G**).

**Figure S3. Time courses of GTPase activity upon opto-GTPase activation.** Supplementary Figure for Figure 2. Time courses of RhoA (**A**), Rac1 (**B**), Cdc42 (**C**), HRas (**D**), Rap1A (**E**), and RalB (**F**) activity for all opto-GTPases were merged into graphs. Time courses for opto-control and opto-GTPases corresponding to GTPase biosensors are the same data presented in Figure 2. Data are presented as the mean ± SD.

**Figure S4. Opto-Ras induced ERK translocation to the nucleus.** HeLa cells transiently expressing opto-GTPases and ERK2-mCherry were serum starved for 6–9 h and observed using a confocal microscope in Leibovitz’s L-15 medium. Opto-GTPases were activated with the help of multi-argon 458-nm laser irradiation every 10 s during a period of 60–350 s. Time course of the relative fluorescence intensity of nuclear ERK2-mCherry (**A**). Changes of the fluorescence intensity of nuclear ERK2-mCherry after 5 min of blue light-irradiation (**B**). Data presented as individual points and the mean ± SD. N = 19, 20, 13, 18, and 11 cells, respectively. Data were analyzed using one-way ANOVA followed by Dunnett’s test between opto-control and other groups. ANOVA *F* = 18.32 and *p* < 0.0001. ***,, *p* < 0.001; ns, not significant; Dunett’s test.

**Figure S5. Intracellular calcium changes in HeLa, MDCK, and HEK293T cells during RhoA activation.** Representative images of HeLa, MDCK, and HEK293T cells expressing opto-RhoA and R-GECO1 in RPE1 cells. From time 0 s, opto-RhoA was activated by a multi-argon 458-nm laser every 10 s. Scale bar, 50 µm.

**Figure S6. Expression of msPLCε-ECFP rescued opto-RhoA-induced calcium transients in PLCε-depleted cells.** siRNA-mediated PLCε-depleted RPE1 and HeLa cells transiently expressing ECFP or msPLCε-ECFP and R-GECO1 were observed (see Figure 3). Data are presented as means ± SD from three independent experiments. In total, >200 cells were analyzed for each condition. **, *p* < 0.01; *, *p* <0.05; two-tailed unpaired Student’s t-test.

**Supplementary Movie Legends**

**Movie 1. Opto-RhoA can control RhoA activity both in time and space.** Supplemental movie for Figure 1C.

**Movie 2. Opto-RhoA induces the calcium transients in various cell types.** Supplemental movie for Figure 3 and S5.

**Movie 3. Opto-5-ptase activation decrease membrane PI(4,5)P_2_.** Supplemental movie for Figure 6E.

**Movie 4. Opto-RhoA activation marginally affects membrane PI(4,5)P_2_.** Supplemental movie for Figure 6E.

**Movie 5. Opto-RhoA-mediated calcium transients translocate NFAT from cytoplasm to the nucleus.** Supplemental movie for Figure 7A.
