## Supplementary figures and images for "Optogenetic control of small GTPases reveals RhoA-mediated intracellular calcium signaling"

### Supplementary Figure S1

**Inaba et al., Figure S1**

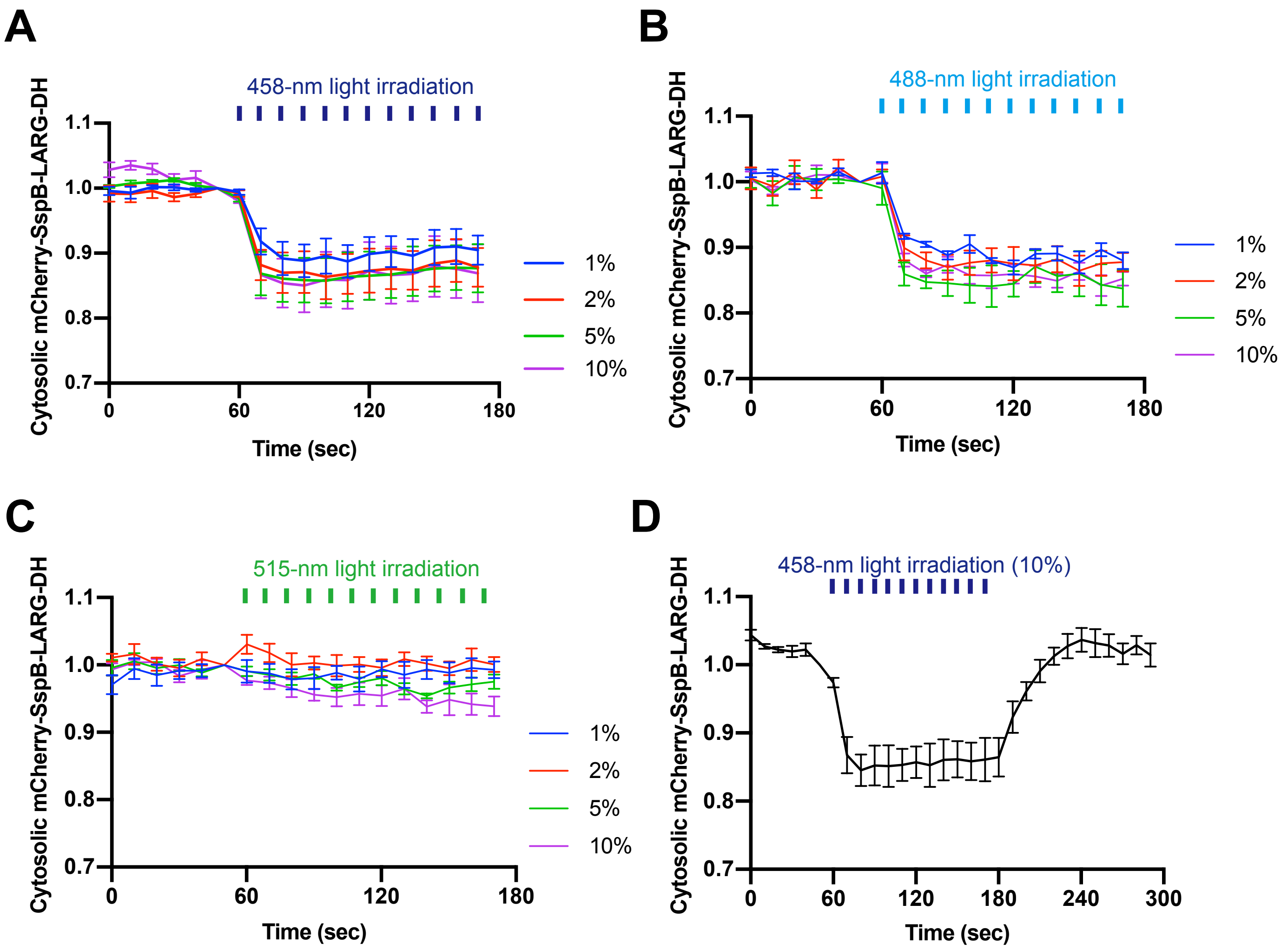

### Supplementary Figure S2

#### Inaba et al., Figure S2

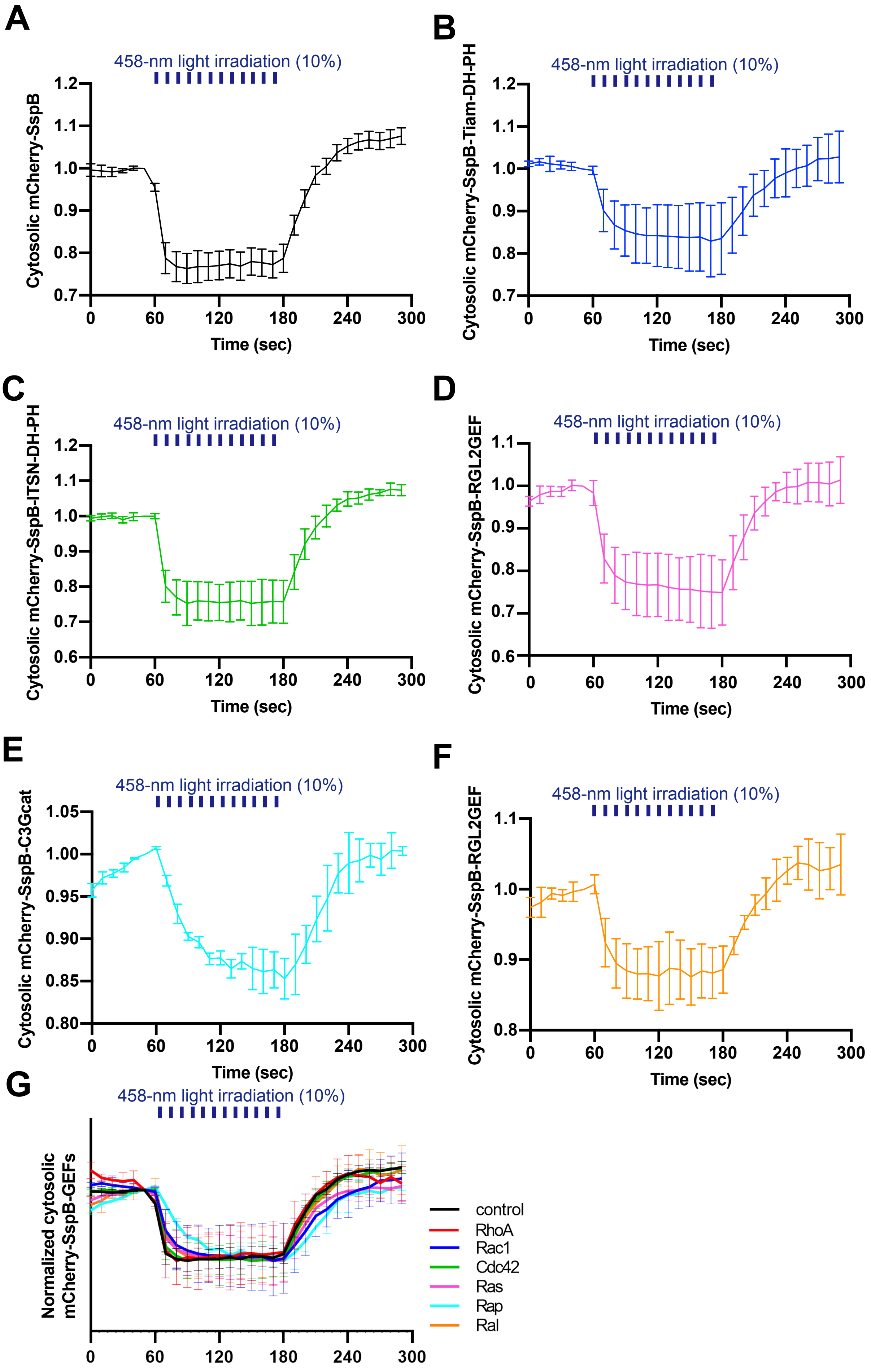

### Supplementary Figure S3

##### Inaba et al., Figure S3

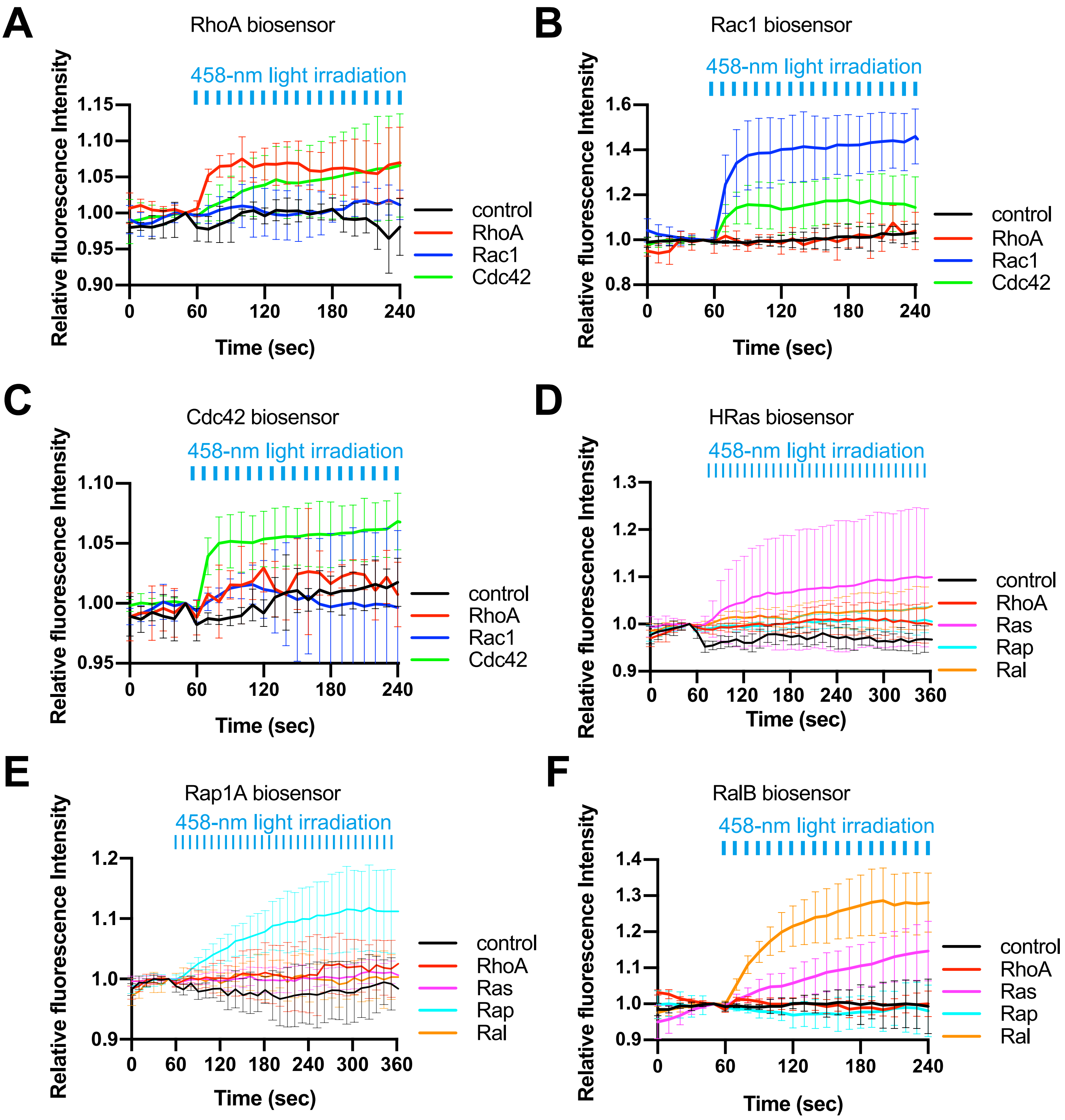

### Supplementary Figure S4

#### Inaba et al., Figure S4

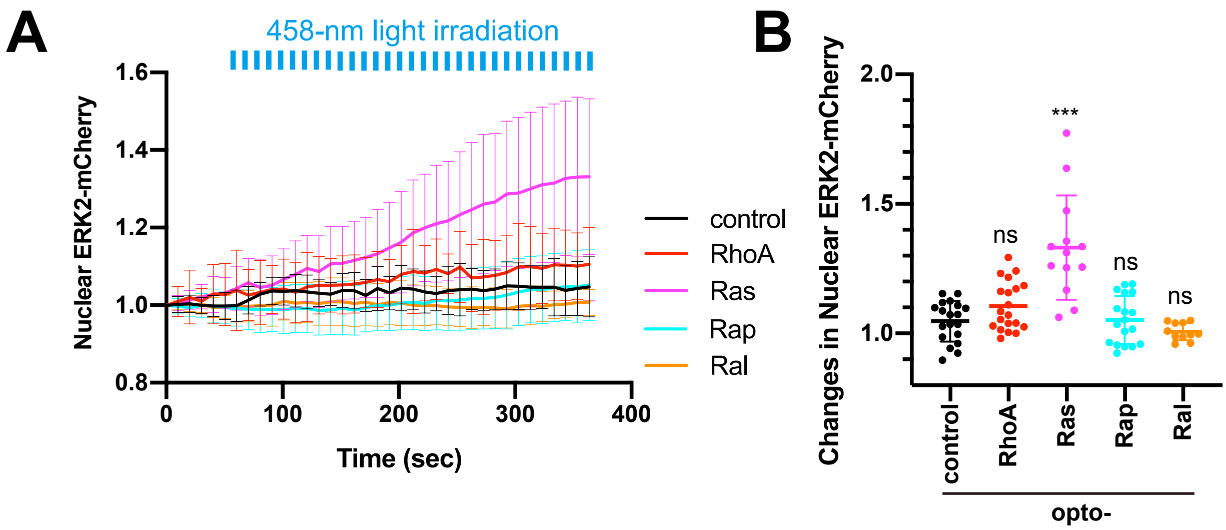

### Supplementary Figure S5

#### Inaba et al., Figure S5

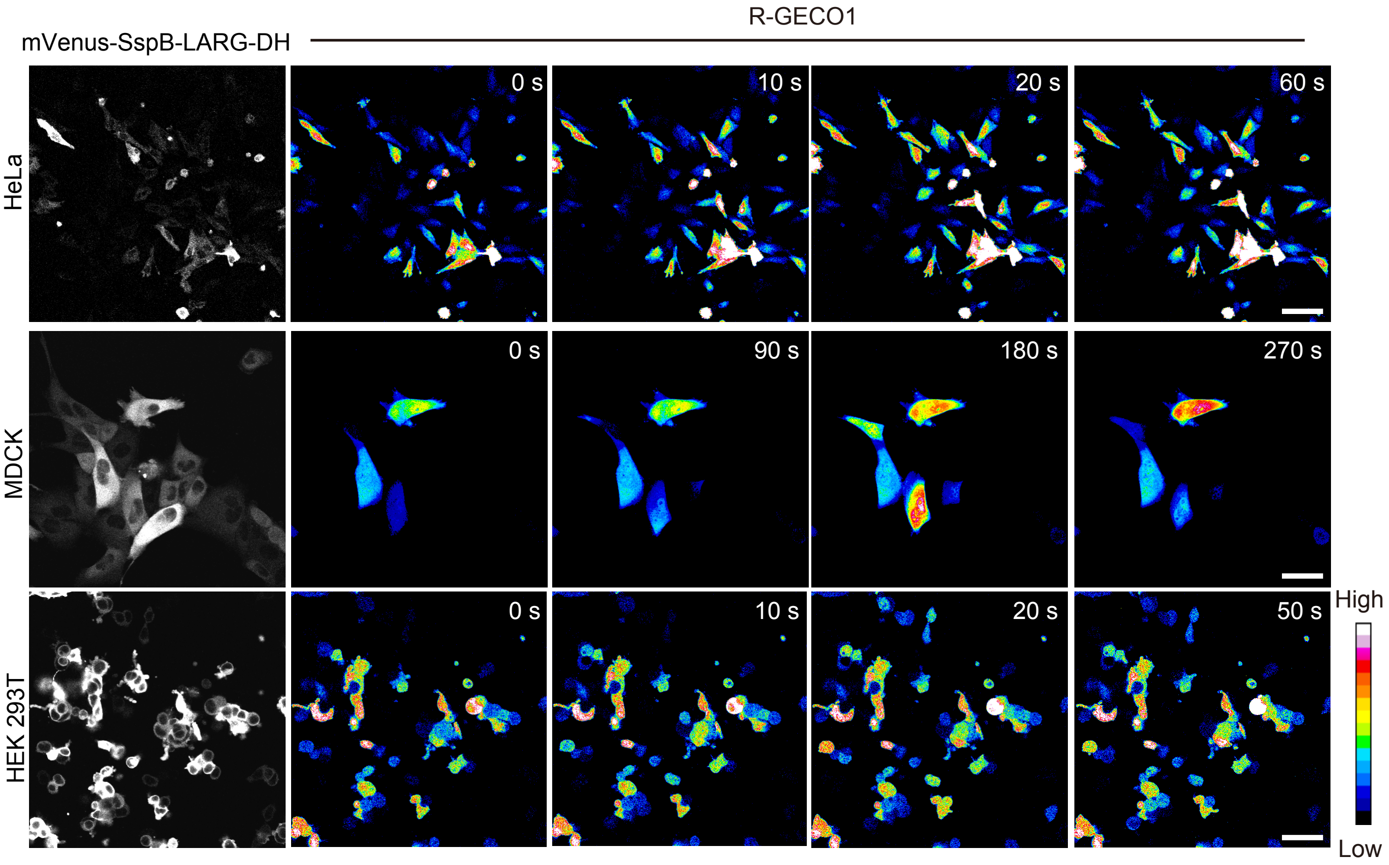

### Supplementary Figure S6

#### Inaba et al., Figure S6

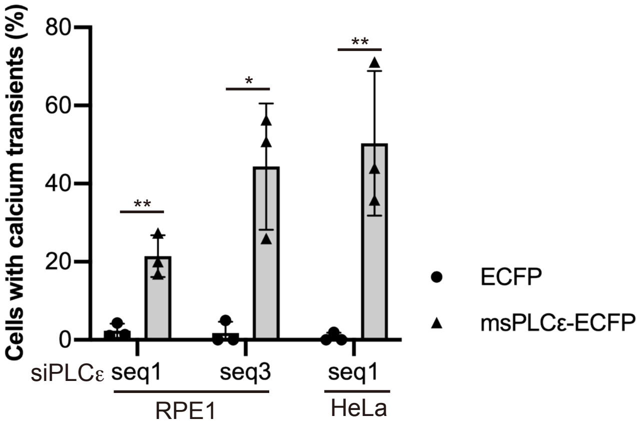
